# Recovery of neural dynamics criticality in personalized whole brain models of stroke

**DOI:** 10.1101/2020.12.17.423349

**Authors:** Rodrigo P. Rocha, Loren Koçillari, Samir Suweis, Michele De Filippo De Grazia, Michel Thiebaut de Schotten, Marco Zorzi, Maurizio Corbetta

## Abstract

The critical brain hypothesis states that biological neuronal networks, because of their structural and functional architecture, work near phase transitions for optimal response to internal and external inputs. Criticality thus provides optimal function and behavioral capabilities. We test this hypothesis by examining the influence of brain injury (strokes) on the criticality of neural dynamics estimated at the level of single participants using directly measured individual structural connectomes and whole-brain models. Lesions engender a sub-critical state that recovers over time in parallel with behavior. The improvement of criticality is associated with the re-modeling of specific white matter connections. We show that personalized whole-brain dynamical models poised at criticality track neural dynamics, alteration post-stroke, and behavior at the level of single participants.

## Introduction

The fundamental mechanisms underlying the dynamics of brain activity are still largely unknown. Interdisciplinary research in neuroscience, inspired by statistical physics, has suggested that healthy brain’s neural dynamics stay close to a critical state^1^, i.e., in the vicinity of a critical phase transition between order and disorder^2,3^, or between asynchronous or synchronous oscillatory activity^4,5^. In physics, critical phenomena occur at the transition of different states of the systems (also known as phase transitions) for specific values of the so-called system’s control parameter (e.g., temperature). There is mounting evidence that biological systems (or parts, aspects, or groups) operate near/at critical points^6,7^. Examples include gene expression patterns^8^, bacterial clustering^9^, flock dynamics^10^, as well as spontaneous brain activity. Indeed, neural systems seem to display features that are characteristic of systems at criticality. These include i) the scaling invariance of neural avalanches^5,11^ reported in diverse species^12,13^, through different imaging techniques^14^, and electro-physiological signals^15^; ii) the presence of long-range spatiotemporal correlations in the amplitude fluctuations of neural oscillations^16,17^, including the observation of 1*/ f* power spectra from simultaneously recorded MEG/EEG signals^15^, fMRI^18^, and cognitive responses^19^; and iii) the increase of the correlation length with system size^17,20,21^.

Critical brains benefit from these emergent features to promptly react to external stimuli to maximize information transmission^22^, hence sensitivity to sensory stimuli, storage of information^23^, and a coordinated global behavior^11,24^. If criticality is indeed a fundamental property of healthy brains^2^, then neurological dysfunctions shall alter this optimal dynamical configuration. However, we know little about the effect of brain disorders on criticality^25^. Some studies have reported disrupted criticality during epileptic seizures^26,27^, slow-wave sleep^28^, anesthesia^29^, sustained wakefulness^30^, states of (un)consciousness^31,32^, and Alzheimer’s disease^33^. However, a crucial test of the hypothesis requires showing alterations of criticality after focal brain injuries that cause local alterations of the brain’s structural and functional architecture. If criticality is essential for behavior, then its alteration after focal injury shall relate to behavioral dysfunction. Over time as behavior improves in the course of recovery, so shall criticality. Finally, we hypothesize that changes in criticality with recovery will depend on specific plasticity mechanisms or functional remodeling as shown in previous fMRI studies^34–37^.

There are several aspects of our investigations. First, we employ a stochastic whole-brain model to simulate large-scale neural dynamics^38^ using as input the directly measured structural connectivity of a stroke patient or healthy control. Importantly, we do not fit the resulting dynamics with empirical measured functional connectivity. The structural connectivity, measured at two time-points: 3 months after stroke (*t*_1_) and one year after stroke (*t*_2_), or three months apart in healthy controls, was used to build personalized whole-brain models. Lesions certainly produces departures from normal structural connectivity, but these structural alterations do not necessarily correspond to an alteration of criticality. Critical dynamics results from the combination of a topology (determined by the structural connectivity) and a given value of excitability. Our method allows measuring departures from criticality at the group level or in individual participants, as well as the recovery of criticality over time.

This approach contrasts with other studies that used average or atlas-based structural connectivity models to simulate activity time courses^39–47^. For example, a recent study by Haimovici et al. found that lesions push the system out of criticality towards a sub-critical state^48^. However, these theoretical results were not validated with real patient data. Other studies found abnormal global metrics of network function, such as information capacity, integration, and entropy in stroke patients as compared to healthy participants^49,50^. However, these models were not personalized, i.e., did not use directly measured individual structural connectivity but healthy group average structural connectomes that were fit with many (hundreds) free model parameters to minimize the distance between the model and the empirical functional connectivity^49^.

Secondly, we apply this computation model strategy to a unique cohort of stroke patients studied prospectively and longitudinally at Washington University in St. Louis. This cohort has been investigated with a large battery of neurobehavioral tests and structural-functional magnetic resonance imaging at 2 weeks, 3 months, and 12 months after the stroke. This cohort is representative of the stroke population both in terms of behavioral deficits, their recovery, and the lesion load location^51^. In previous work, we have characterized the behavioral, structural, and functional connectivity abnormalities in this cohort and their relationship to behavioral impairment and recovery^34–37,52^ (see Corbetta et al.^53^ for a review). In network terminology, strokes cause an acute decrease of modularity that normalizes over time^34,54^.

In this work we use stroke as the prototypical pathological model of human focal brain injury and whole-brain computational models to estimate neural dynamics, related alterations in criticality and behavior, and the underlying neural mechanisms. We show that stroke lesions engender a sub-critical state characterized by decreased levels of neural activity, entropy, and functional connectivity, that recovers over time in parallel with behavior. The improvement of criticality is associated with specific white matter connections remodeling.

## Results

### Simulation of large-scale neural dynamics

To simulate neural activity at the individual whole-brain level we employed the homeostatic plasticity model recently developed in^21,38^. Figure 1 illustrates the main ingredients of our modeling strategy. Individual structural connectivity matrices are the key inputs of the stochastic model (Fig. 1A). Imaging and behavioral data are taken from a large-scale stroke study described in previous publications^35,51,55^. Structural connectivity data was available for 79 patients, acquired three months (*t*_1_) and one year (*t*_2_) after stroke onset. The same study includes data from 28 healthy controls, acquired twice three months apart. The difference in the time interval between stroke and controls is that diffusion imaging in patients was obtained only at three and twelve months since the diffusion signal is highly variable early post-injury. In contrast behavioral, structural, and functional connectivity were obtained at all three time points in both groups (see Methods sections for details about the stroke dataset, lesion analysis, diffusion weighted imaging (DWI), and resting-state Functional magnetic resonance imaging).

**Figure 1.**
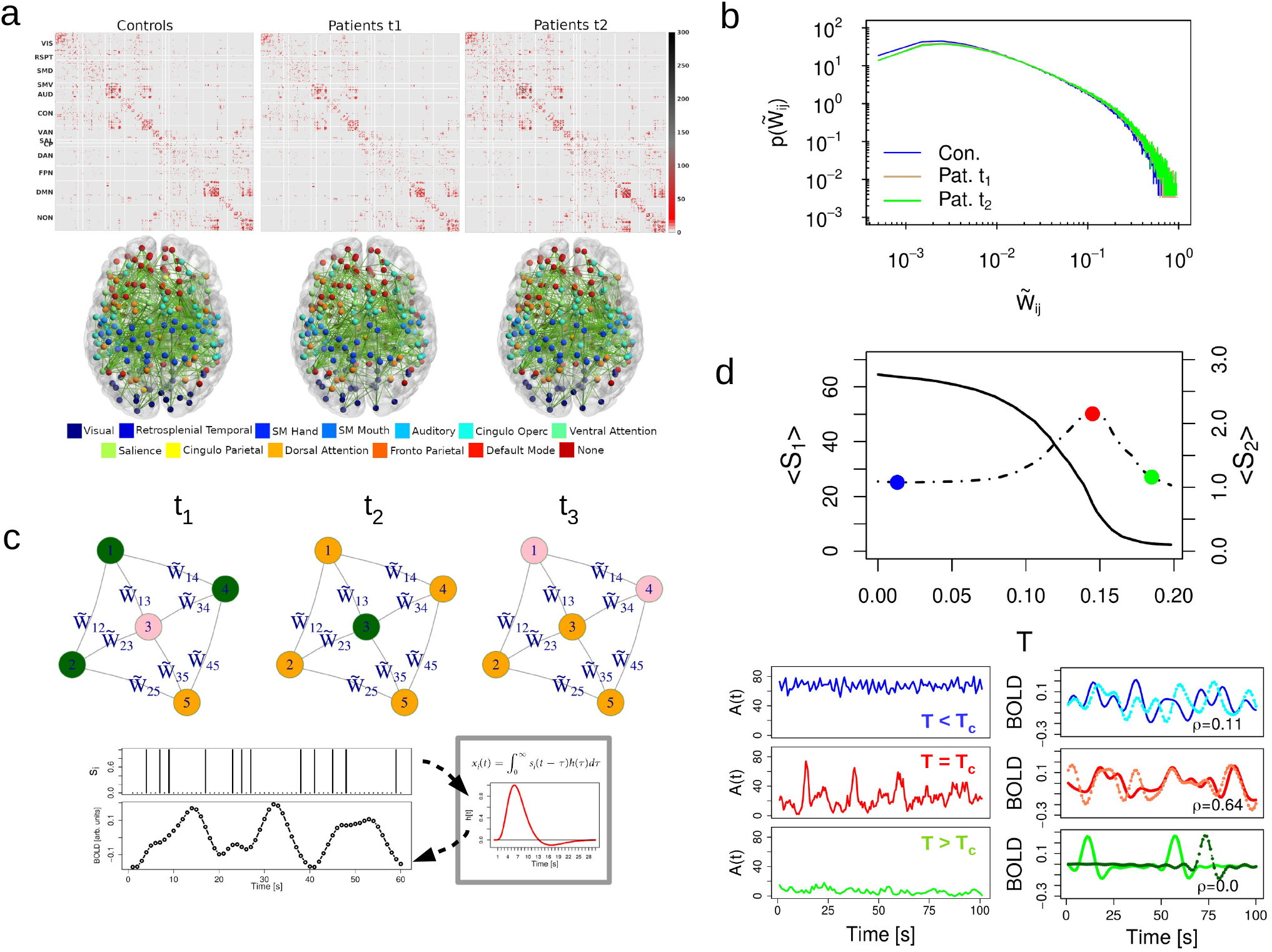
Overview of whole-brain modeling. a) Average structural connectivity (SC) matrices (top) and their corresponding network architecture embedded in a glass dorsal view of the brain (bottom). SC matrices and brain networks are organized according to regions of interest (ROI) defined on the cortical parcellation of Gordon et al.^59^. b) The probability distribution function of the structural connectivity weights in controls and patients after homeostatic normalization (see main text, Eq. (1)). For each group, the (non-zero) weights of all individual matrices were pooled together and then the histogram was computed leading to a representative pdf for the corresponding group. c) Top. Illustration of the network dynamics with homeostatic plasticity following the transition probabilities between the three possible states: inactive (*I*), active (*A*) and refractory (*R*). The temporal evolution of the central inactive node (pink) is as follows: in *t*_1_, it is surrounded by three active nodes (green) and one refractory node (orange); in *t*_2_, the incoming excitation is propagated 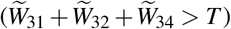; and finally, in *t*_3_, it reaches the refractory state. Bottom. Procedure used to transform node’s activity, *s*_*i*_(*t*), in functional BOLD signals, *x*_*i*_(*t*). BOLD time-series are obtained by convolving instantaneous *s*_*i*_(*t*) with a canonical hemodynamic response function (HRF). d) Behavior of the neural variables, the largest (*S*_1_, continuous line), and the second largest (*S*_2_, dotted line) cluster size as a function of *T* (top). The peak in *S*_2_ (red dot) is identified as the critical phase transition^62^. Blue and green dots correspond to minimal and maximal values of *T*, and corresponding activity and BOLD time-series in the lower panels. Left panel: instantaneous network activity, *A*(*t*) = ∑_*i*_ *s*_*i*_(*t*), for different values of the activation threshold *T* ; the super-critical phase *T* ≪ *T*_*c*_ (blue time-series), the critical phase *T* = *T*_*c*_ (red) and the sub-critical phase *T* ≫ *T*_*c*_ (green). Right panel: example of the simulated BOLD signals between two arbitrary ROIs and their corresponding Pearson correlation *ρ*. The highestcorrelation is achieved at the critical phase, where BOLD fluctuations are long-range correlated.

The whole-brain is described as a network of *N* = 324 nodes (i.e., cortical brain regions), linked with symmetric and weighted connections obtained from DWI scans and reconstructed with spherical deconvolution^56,57^. The weights of the structural connectivity matrix, *W*_*i j*_, describe the connection density, i.e, the number of white matter fiber tracts connecting a given pair of regions of interest (ROIs) normalized by the product of their average surface and average fiber length^58^. The ROIs are derived from a functional atlas of the cerebral cortex^59^. Fig 1A shows the topography of structural connections, the corresponding network assignment, and the corresponding averaged structural matrices. The matrix is sparse, and contains many short-range connections and fewer long-range connections. In the control group, inter-hemispheric connections between homotopic regions of the same network are visible (dorsal view of the brain Fig. 1A). In stroke patients, inter-hemispheric connectivity is decreased, as shown by Griffis et al.^35^ who found that loss of inter-hemispheric connections - both structural and functional - is the predominant aberrant pattern in stroke. The lesion load topography in the stroke cohort is shown in Supplementary Figure 1 and matches the topography of larger cohorts^51,60^.

Cortical activity is modeled through stochastic dynamics based on a discrete cellular automaton with three states, namely, active (*A*), inactive (*I*), and refractory (*R*). The state variable of a given node *i, s*_*i*_(*t*), is set to 1 if the node is active and 0 otherwise. The temporal dynamics of the *i*-th node is governed by the following transition probabilities between pair of states: (i) *I* → *A* either with a fixed small probability *r*_1_ ∝ *N*^−1^ or with probability 1 if the sum of the connections weights of the active neighbors *j*, ∑ _*j*_ *W*_*i j*_, is greater than a given threshold *T*, i.e., ∑ _*j*_*W*_*ij*_*s*_*j*_ > *T*, otherwise *I* → *I*, (ii) *A* → *R* with probability 1, and (iii) *R* → *I* with a fixed probability *r*_2_^38^. The state of each node is overwritten only after the whole network is updated. Therefore, during the temporal dynamics, a node activation happens (most frequently) when the incoming input excitation from its nearest active neighbors exceeds a fixed threshold *T*, i.e, ∑ _*j*_*W*_*ij*_*s*_*j*_ > *T*. In other words, *T* plays the role of a threshold parameter that regulates the propagation of incoming excitatory activity. On the other hand, the two parameters *r*_1_ and *r*_2_ control the time scale of self-activation and recovery to the excited state^21,38^. As we clarify in the methods section, *r*_1_ and *r*_2_ are set as a function of the network size, while *T* is the control parameter of the model.

Following^38,61^, we consider homeostatic plasticity principles regulating network excitability by introducing a normalization of the structural connectivity matrix

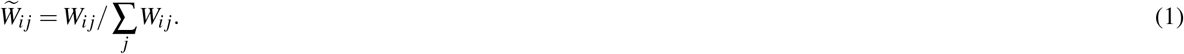

As shown by^38^, the above normalization minimizes the variability of the neural activity patterns and the critical point of the stochastic model for different participants, facilitating statistical comparison among model outputs for single individuals. Moreover it facilitates the emergence of functional networks at rest, and increases the correlation coefficients between model and empirical data.

For each participant (stroke, control) and time point (*t*_1_ : 3 months for stroke, first scan for control; *t*_2_ : 12 months for stroke, second scan three months apart for control) we calculate the following neural state variables (see Methods section): the average activity (*A*), the standard deviation of the activity (***σ***_*A*_), and the size of the averaged clusters, the largest (*S*_1_), and the second largest (*S*_2_), as a function of the activation threshold *T*. These clusters of activity are defined as the size of the connected components of the graph defined by the sets of nodes that are both structurally connected to each other and simultaneously active^38^. In the numerical experiments, we set the total simulation time-steps *t*_*s*_ = 2000, to approximate the length of a typical fMRI experimental time-series (∼ 15 min).

A typical behavior of the simulated brain activity for different values of *T*, while keeping *r*_1_ and *r*_2_ fixed, is illustrated in Fig. 1C. The brain dynamics displays a phase transition at critical threshold *T*_*c*_ given by the corresponding value of *T*. In fact, at *T* = *T*_*c*_ brain activity has the largest variability, the maximal second largest cluster size, and a steep change in the first cluster size^21,38,62^. In contrast, for small values of the activation threshold (*T* ≪ *T*_*c*_), the system is characterized by high levels of excitation, i.e, the signal from an active node will easily spread to its neighbors. In this scenario, we have the so-called super-critical or disordered phase, which is characterized by sustained spontaneous activity with fast and temporally uncorrelated fluctuations (Fig. 1D, blue time-series). On the other hand, high values of *T* (*T* ≫ *T*_*c*_) lead to a sub-critical or ordered phase, which is characterized by regular, short propagating and not self-sustained activity. In this case, only those nodes with the strongest connections will determine the excitation flow in the network. In the sub-critical phase, simulated BOLD signals have very small correlations (Fig. 1D, green time-series). The critical phase appears in between of these two states, when brain activity displays oscillatory behavior and long-range temporal correlations of their envelope^21,38,48^. At criticality the simulated BOLD activity shows the highest correlation (Fig. 1D, red time-series)^38^.

In the next sections, first, we present simulations of the whole brain model with homeostatic plasticity for stroke and control individuals, as well as at the group level. Next, we study the structural connectivity correlates of neural dynamics alterations induced by stroke lesions. Third, we identify components of the structural networks that are most strongly related to criticality and its recovery over time. Fourth, we compare the simulations output with empirical functional networks, and with behavioral data obtained in multiple domains (e.g., language, motor, memory) from an extensive neuropsychological battery (see Methods section and Supplementary Note 2).

### Abnormal neural dynamics in stroke

In this section we shall investigate the model’s fingerprint of criticality loss. Stroke is not a binary phenomenon, thus we do not expect that all individuals will lose criticality following a lesion. While some patients neural patterns will behave similarly to controls others will significantly depart from normality. First, we show the fingerprints of criticality loss and associated recovery in a representative patient, then we quantify the variability of the neural activity patterns across groups and time points. All the statistical tests reported throughout this study met the assumptions of normality and equal variances.

Figure 2(A-D) shows the model’s neural activity variables for an arbitrary control-patient pair. The neural profiles for each stroke patient and healthy control can be seen in the Supplementary Figures 10-16. To facilitate the comparison between individual control and stroke data, we also present the average and standard deviation of the healthy controls (blue dashed line and shaded area, respectively). In general, the neural dynamics in healthy participants are quite distinct from stroke patients who manifests a loss of criticality. First, let us consider one healthy participant, Fig. 2(A-B) (black/gray dots). The neural patterns follow the expected behavior, with a critical point *T* = *T*_*c*_ around the maximum of *S*_2_, or equivalently, near the sharp decrease of *S*_1_. Moreover, as expected, the two curves have low variability across the two time points (*t*_1_ and *t*_2_), displaying the same *T*_*c*_ (within one standard deviation), and stable shape as a function of *T*.

**Figure 2.**
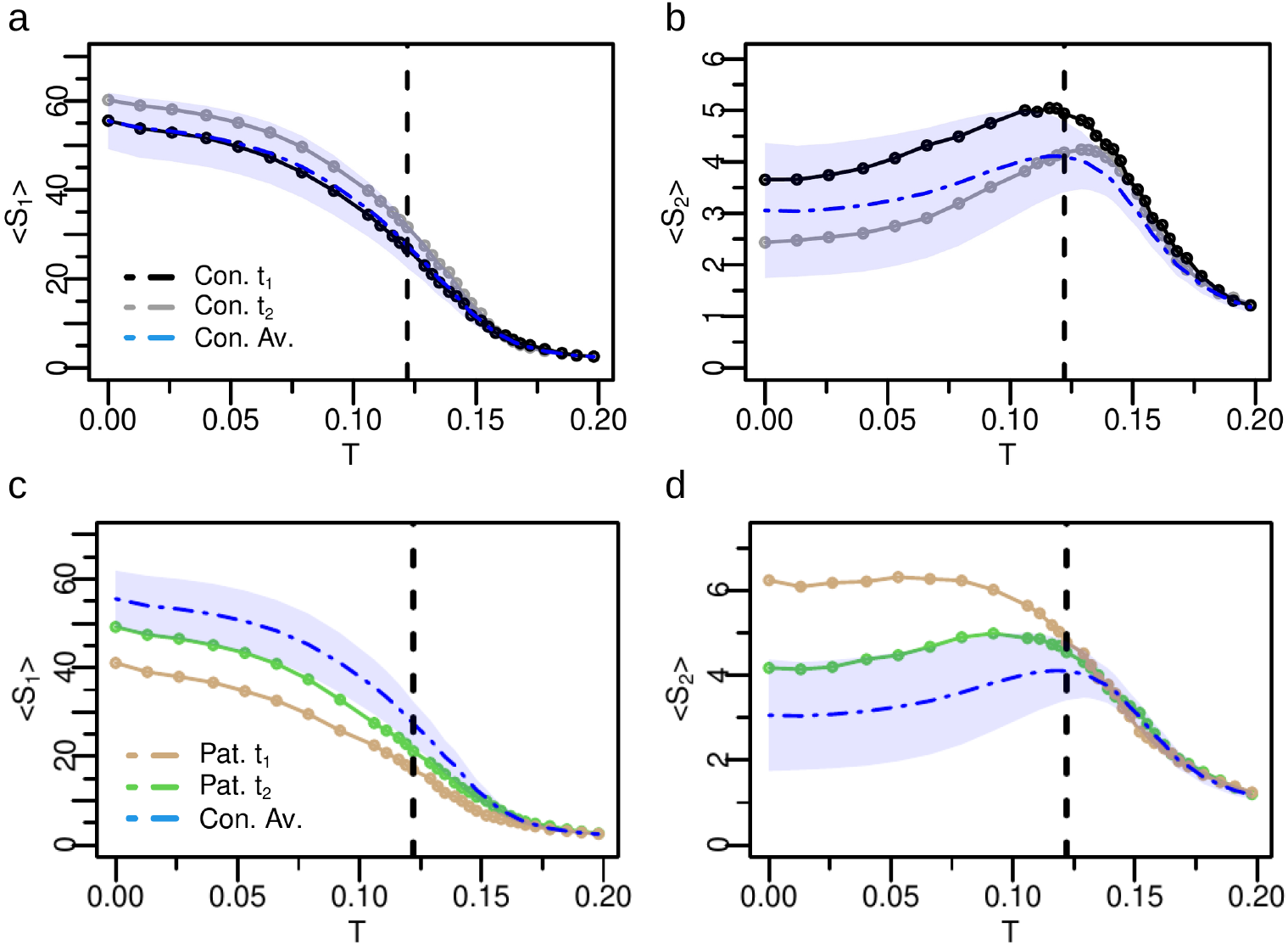
Fingerprints of criticality loss and associated recovery in a representative patient. Analysis of neural activity patterns, *S*_1_ and *S*_2_, of a healthy participant (Con. / a-b) and stroke patient (Pat. / c-d). In blue dashed line we show the corresponding control’s group average, while the shaded area corresponds to one standard deviation. For the healthy participant, *t*_1_ and *t*_2_ correspond at two different time points 3 months apart. For the patient, *t*_1_ and *t*_2_ correspond to 3 months and 12 months post-stroke. The healthy participant exhibits features of criticality at both time points with small variability across time points and within the variability of the healthy group. At the same threshold *T*, both *S*_1_ and *S*_2_ sharply change their size. For the stroke patient under consideration, the flattened shape at *t*_1_ of both, *S*_1_ and *S*_2_, combined with the monotonic behavior of *S*_2_ indicates lack of criticality, which, however, returns at *t*_2_ as depicted by the emergence of a peak in *S*_2_. The black vertical dashed line depicts the critical point of healthy controls. Source data are provided as a Source Data file.

The pattern is dramatically different in the stroke patient. The characteristic peak in *S*_2_^38,62^ describing the critical phase transition is completely absent at three months post-stroke, but normalizes at one year, where the transition is sharper and both, *S*_1_ and *S*_2_, with a behavior similar to normal. The monotonic behavior of *S*_2_ is associated with a loss of a critical phase transition (for any value of excitability *T*), as shown in a recent study by Zarepour and colleagues^62^. Accordingly, this monotonic behaviour is encountered when the average degree or the disorder of the structural connectivity matrix 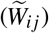 is low. When the connectivity disorder or degree increases then criticality emerges. Therefore, the recovery of criticality at 12 months predicts a reorganization of the topology of the structural connectome, with an associated increase of both, the average degree and the connectivity disorder, as a function of time. We shall investigate the anatomical bases of brain criticality modifications in the next section.

As already observed in^38^, the homeostatic normalization on the weights of the structural matrix decreases the inter-participant variability of neural activity patterns. More importantly, it fixes the critical point of healthy controls to a universal value, *T*_*c*_ ∼ 0.122. In other words, thanks to the homeostatic plasticity mechanism, the critical point is independent of the individual variability in the structural connectivity matrix. However, the strength of the *S*_1_ and *S*_2_ peaks provides the characterization of differences in criticality. In fact, near the critical point the differences in the stroke patient at *t*_1_ and *t*_2_ are pronounced. The most interesting feature, as shown in Figs. 2(C-D), is the recovery-like pattern: the one-year post-stroke curve resembles, both qualitatively and quantitatively, the pattern of the healthy controls. Similar results are shown for several examples of individual stroke patients and healthy participants (see Supplementary Figures 10-16 and Supplementary Note 4).

We next examined the variability of the neural activity patterns across groups and time points. Figure 3 summarizes the results for *S*_1_, *S*_2_, *A* (the average activity) and *σ*_*A*_ (the variability of activity), respectively (the functional connectivity (FC) and the entropy (*H*) are shown in Supplementary Figure 3). The thin solid curves represent each individual stroke patient, while the thick dashed lines (brown = stroke 3 months; green = stroke 12 months; blue = average control) represent the group average, i.e., 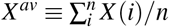, where *n* is the number of individuals in each group. Individual stroke patients, as expected, showed great variability of *S*_1_, *S*_2_ behavior at *t*_1_ with some exhibiting loss of criticality (monotonic *S*_2_ and flat *S*_1_) while others showing normal curves. On average, however, the stroke group at *t*_1_ manifested a less critical behavior as apparent both in the distribution of individual (brown) curves and the average curves. In *t*_2_, this abnormal condition was restored, and *S*_2_ decreased toward normal, while *S*_1_ increased. These changes in cluster sizes reflect alterations in segregation-integration balance within/between networks^63^. The total average activity, *A*, followed the behavior of *S*_1_. The variability of neural activity, *σ*_*A*_, presented a sharp peak at the critical point for healthy participants, but was significantly attenuated in patients. We statistically assessed how the neural variables evaluated at the critical point shown in Fig. 3 varied between groups and across the two time points using mixed ANOVA. The group main effect was reliable for all variables (all p-values < 0.004), confirming the alterations in the neural dynamics for stroke patients with respect to healthy controls. However, the main effect of time was not significant. We expected a group by time interaction reflecting longitudinal changes restricted to the patient group, but this was only significant for the standard deviation (p-value = 0.022) due to the large variability across patients. Nevertheless, we carried out planned paired two-tailed t-tests to further assess changes in the patients’ neural variables over time. We found statistically significant differences for *S*_1_ (p-value = 0.03) but not for *S*_2_ and *A* (Supplementary Table 3).

**Figure 3.**
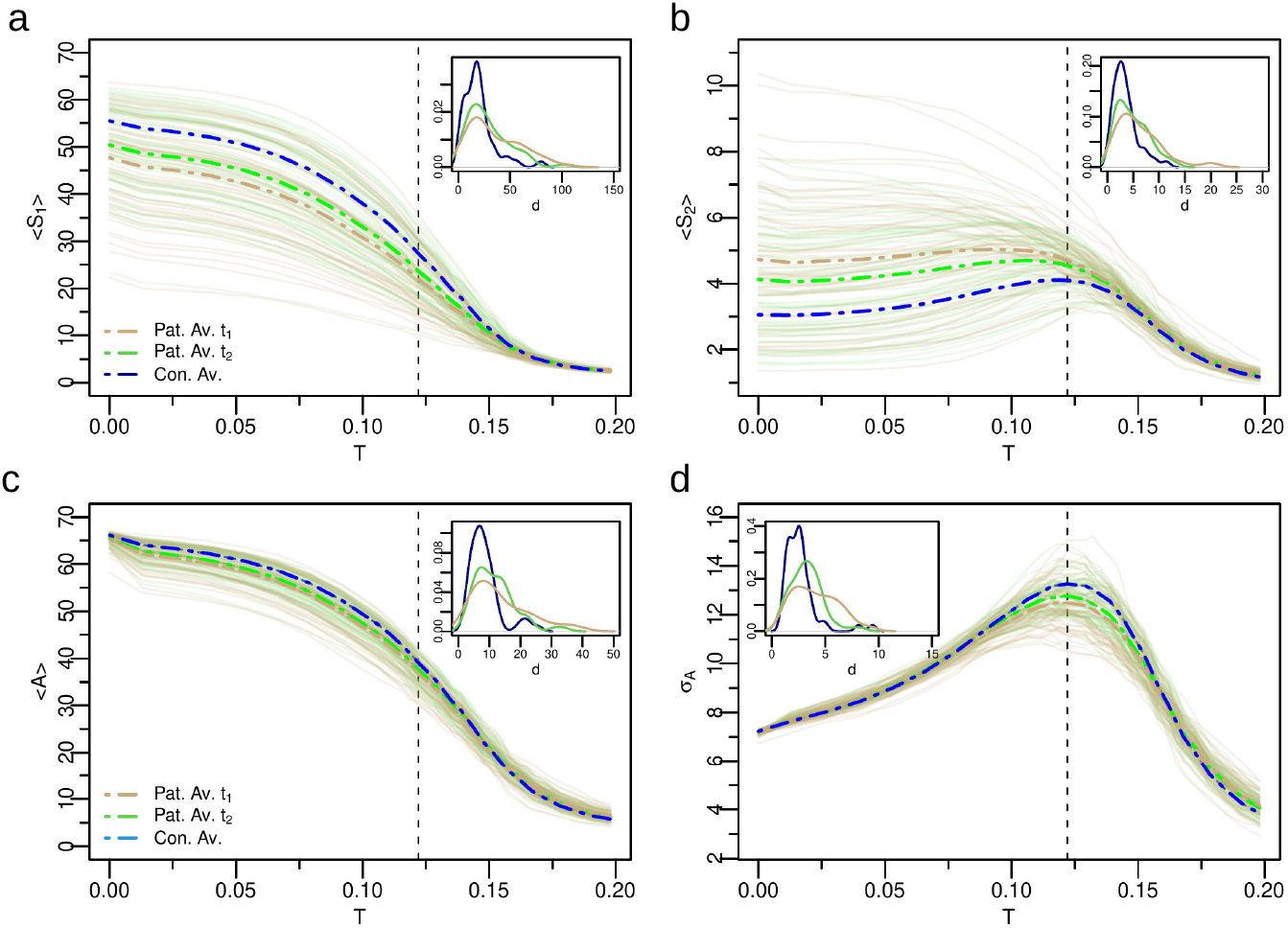
Variability of the neural dynamics criticality across groups and time points. a-b) Group based analysis of neural activity patterns, *S*_1_ and *S*_2_, as a function of *T* for all patients and controls. The brown lines represent patients at *t*_1_ (3 months post-stroke, *n* = 54), and the green lines at *t*_2_ (12 months post-stroke, *n* = 59). The thin solid curves indicate each individual stroke patient, while the thick dashed lines represent the group average. The group analysis reveals that on average patient’s neural activity patterns are abnormal at *t*_1_, but approach control levels at *t*_2_. c-d) Average activity and standard deviation of activity, respectively. Note improvement of the activity and its variability across time points. Insets: Probability distribution function (pdf) of the Euclidean distances *d* in individual age-matched-controls and stroke patients at *t*_1_ (brown) and *t*_2_ (green). Patients at *t*_1_ show greater variability of model neural activity as depicted by the longer pdf’s tails. Note trend toward normalization from *t*_1_ to *t*_2_. The black vertical dashed line depicts the critical point of healthy controls. Source data are provided as a Source Data file.

The Euclidean distance, *d*, was then used to quantify the similarity between individual’s neural activity pattern (i.e, *X* (*i*)) with the corresponding control group average (see Methods section (9)). This measure takes in consideration the behavior of the neural profile across all values of excitability *T*. A low *d* indicates low variability of the dynamic parameters across participants and time points as observed in health participants (e.g. Figure 2 A-B). A high *d* indicates high variability across participants and time points of the neural state variables implying the presence in the group of abnormal dynamics. The insets in figure 3 show the normalized distribution of the Euclidean distances for each variable (*S*_1_, *S*_2_, *A, σ*_*A*_). Each variable shows higher variability in stroke than controls, and higher variability at *t*_1_ than *t*_2_. Based on^62^ changes in criticality of the model must depend on changes in the underlying structural connectivity, a biological prediction that we will examine in more detail in the next section.

In summary, both at the level of single participants and group level, the models of stroke patients show a significant loss of the critical dynamics at three months that recover on average at one year. This is consistent with the first hypothesis that criticality is a property of the normal brain structural architecture. Next, we examine the anatomical bases of brain criticality modifications.

### Structural connectivity explains criticality and its recovery

The recovery of criticality from three to twelve months must reflect a change in the underlying structural connectivity. Therefore, we applied measures from graph theory to quantify the topology of the corresponding structural brain networks and associated alterations following stroke recovery. We choose the average degree as a measure of the overall network connectivity (density), the modularity and the global efficiency as measures of the degree of network segregation and integration^64^, respectively. Furthermore, to quantify the connectivity disorder, we defined the entropy of the homeostatic connectivity matrix (*H*_*SC*_) in an analogous way to the functional connectivity entropy^50^ (see Method’s section).

Inspired by Zarepour et al^62^, we built a map-like parameter space using each participant average degree (*K*) and connectivity disorder (*H*_*SC*_) for *t*_1_ and *t*_2_. We identified stroke patients with *S*_2_ monotonic decay (through numerical derivative, see Method’s section) to obtain the critical value of average degree *K* and connectivity disorder *H*_*SC*_ below which the critical transition disappears (triangular dots in Fig. 4 A). This critical threshold was sharp and close to *K*_*c*_ ∼ 14 and *H*_*c*_ ∼ 0.055 (the healthy participant’s average degree is *K* = 18 ± 3 and connectivity disorder *H*_*SC*_ = 0.059 ± 0.004). Therefore, the change in the criticality regime is associated with a decrease of the average degree and the connectivity disorder below *K*_*c*_ and *H*_*c*_, respectively. Healthy controls and patients at *t*_1_ showed quite separate distributions for *K* as well as *H*_*SC*_, while these values tended to normalize at *t*_2_. Still there were several stroke patients who maintained a loss of criticality at *t*_2_ (green triangles). The recovery of criticality from *t*_1_ to *t*_2_ was associated with an increase of both the average degree (paired t-test: *t* = 2.1, p-value = 0.04) and the connectivity disorder (paired t-test: *t* = 2.2, p-value = 0.03). The amount of recovery can be captured by the recovery indexes *H*_*SC*_(*t*_2_ −*t*_1_) and *K*(*t*_2_ −*t*_1_), defined as the difference of *H*_*SC*_ and *K* between the two time points. We found a significant positive correlation between the latter two quantities (*R*^2^ = 0.87, *ρ* = 0.93 and p-value< 10^−16^, inset of Fig. 4 A). In summary, changes in criticality regime appear to be strictly related to changes of the stroke patient’s network average degree and connectivity disorder in accordance with the theory^62^.

**Figure 4.**
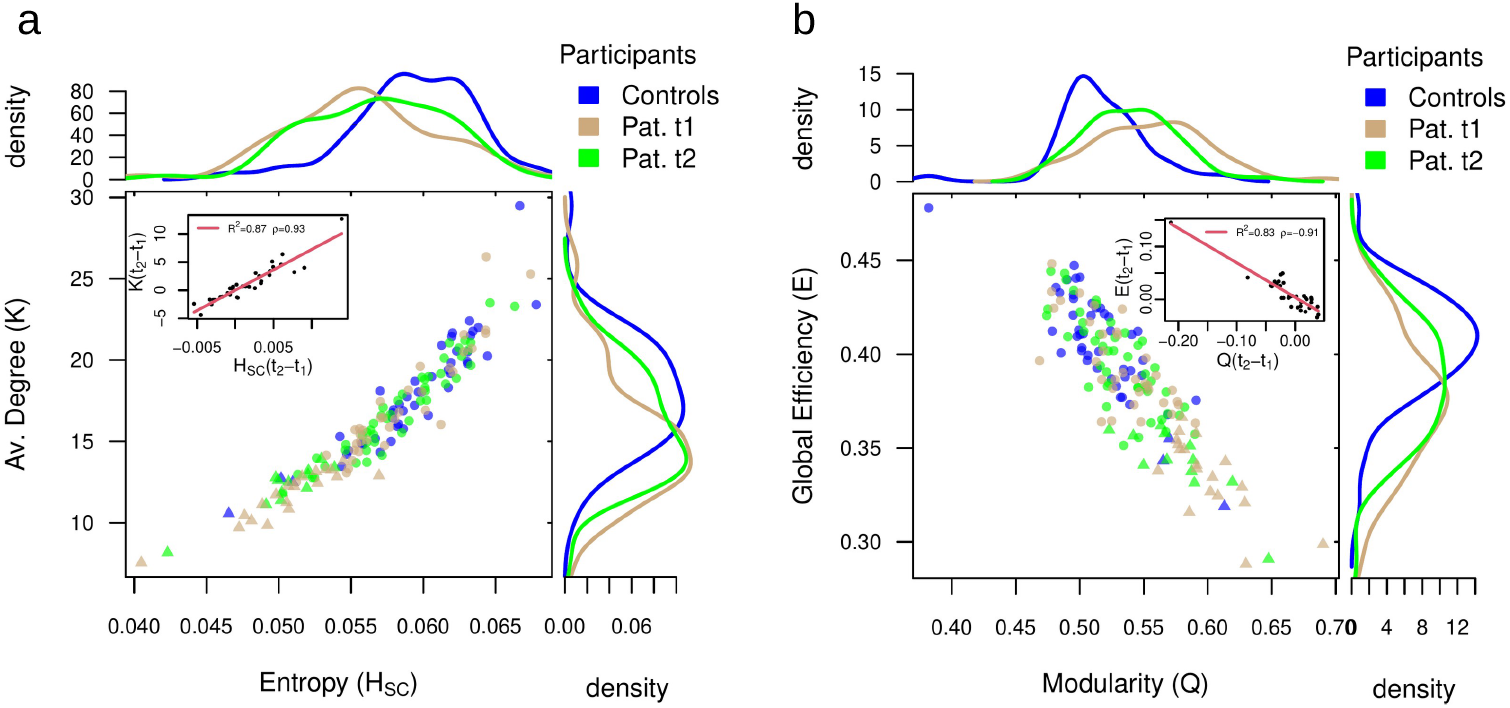
Alterations in network topology and associated changes in criticality regime. a) Average degree (*K*) versus connectivity disorder (i.e., structural entropy *H*_*SC*_) for all patients and controls. Controls are colored in blue (*n* = 46). Brown dots represent patients at *t*_1_ (3 months post-stroke, *n* = 54), while green dots at *t*_2_ (12 months post-stroke, *n* = 59). Normalized density plots (histograms) are also shown. Triangular dots correspond to patients who have *S*_2_ monotonic decay and so lack of criticality. In the inset we show the recovery indexes (see main text) *K*(*t*_2_ − *t*_1_) versus *H*_*SC*_(*t*_2_ − *t*_1_). In the legend we show the (linear) correlation, *ρ* and the *R*^2^. Normalization of both, the average degree and the connectivity disorder, from *t*_1_ to *t*_2_ supports the corresponding recovery of patients criticality. b) The same as a) but for the global efficiency (*E*) and modularity (*Q*). Strokes cause an overall decrease in global efficiency at the cost of an associated increase in modularity, but that recovers from *t*_1_ to *t*_2_ towards the controls. Source data are provided as a Source Data file.

Then we analyzed the effect of lesion size on *K* and *H*_*SC*_, to disentangle its contribution for the criticality regime alterations. The lesion size predicted a small amount of variance for both variables. We find a small negative correlation with the average degree (*t*_1_: *ρ* = −0.26, p-value = 0.051; *t*_2_: *ρ* = −0.15, p-value = 0.25). The lesion size captured slightly more variance at the first time-point, however, the statistical significance was lost after correcting for multiple comparisons. For the connectivity disorder, the correlation with lesion size was even weaker (*t*_1_: *ρ* = −0.21, p-value = 0.11; *t*_2_: *ρ* = −0.09, p-value = 0.5). Furthermore, recovery indexes of both variables did not correlate with lesion size (*ρ* ∼ 0). These results indicate a weak relationship between lesion size and loss of criticality.

We further characterized the brain network organization in terms of modularity (an index of network segregation) and global efficiency (an index of network integration). Healthy participants are highly clustered in the upper left region of the scatter plot and show the highest values of global efficiency and lowest values of modularity. This pattern reflects a balanced network configuration that supports functional segregation between distinct specialized brain regions while allowing for functional integration. However, stroke disrupts this balance. In fact, patients at *t*_1_ show decreased global efficiency reflecting a decrement of network integration with reduced capacity for information transfer between distant brain regions. At *t*_2_ global efficiency and modularity tend to normalize moving their distribution toward controls. Changes of modularity from *t*_1_ to *t*_2_ are negatively correlated with changes in global efficiency (*R*^2^ = 0.8, *ρ* = −0.9 and p-value< 10^−14^, inset of Fig. 4 B). A pathological increase of network segregation therefore comes at the cost of diminished integration and vice-versa. The global efficiency presents more robust variations across time points (paired t-test: *t* = 1.8, p-value = 0.08) than modularity (paired t-test: *t* = −1.2, p-value = 0.23). In summary these analyses indicate remodeling of white matter connections from three to twelve months post-stroke that correspond to changes in network organization. In the Supplementary Information we provide additional evidence for this re-modeling (Supplementary Table 1 and Supplementary Note 1). We tested whether the structural connectivity matrices, or portions of them, changed over time in terms of the number of white-matter fibers. We found significant global changes (averaging across all nodes) from 3 to 12 months both in the number of fibers and their topology. We also found significant increases in the number of fibers at the level of many brain networks. In contrast, no significant differences from *t*_1_ to *t*_2_ were detected in healthy controls.

Next, we investigate which connections were more strongly related to the alteration and recovery of criticality. To this end, we used a multivariate machine learning approach, based on a cross-validated Ridge Regression^65^, to relate the model’s neural activity variables to the structural connectivity matrix. This approach allows to identify edges (and sub-networks) across the whole brain that are most strongly related to the variable of interest (see Methods for details).

First, we investigate the relationship between structural connectivity and criticality, both in healthy participants and in stroke patients. We employ the threshold independent dynamical variable *I*_2_, defined as: *I*_2_ = ∫*S*_2_*dT*. The rationale behind the choice of this variable is twofold. First, *I*_2_ is positively correlated to *S*_2_ evaluated at the critical point (*ρ* ∼ 0.9 and p-value< 10^−16^), but it is much more stable. Indeed, the integration of *S*_2_ over the entire range of excitability smooths out the fluctuations due to the intrinsic variability in brain topologies and associated dynamics. Second, this variable predicts a large amount of behavioral performance at each time point separately (see next section). Figure 5 shows that the structural connectivity accounts for a large proportion of variance in *I*_2_ (controls: *R*^2^ = 0.68; patients at *t*_1_: *R*^2^ = 0.57; patients at *t*_2_: *R*^2^ = 0.69). The map of predictive edges, *W*_*i j*_ (see (14)), plays a key role in understanding the alterations of neural dynamics criticality (*I*_2_) vis-a-vis white-matter re-modeling. We provide two distinct complementary visualizations of *W*_*i j*_. First, we display the top 200 edges embedded in an anatomical space (top, Fig. 5). Small values of *I*_2_ correspond to neural patterns compatible with critical dynamics. Thus, positive edges (colored orange) contributes to criticality loss (*W*_*i j*_ > 0), while negative edges (colored green, *W*_*i j*_ < 0) reinforce the emergence of critical dynamics. In healthy participants predictive edges concentrate in auditory, cingulo-opercular, default mode, fronto-parietal and somato-motor mouth networks; interestingly, the visual network contributes with few regions with weak edges. Healthy participants show a predominance of negative edges (positively associated with the emergence of criticality). In contrast, predictive maps of structural connections in stroke patients show a different organization. The connectivity weights are not as balanced as controls (see width of the edges). The visual network is more prominent in patients, especially at *t*_2_.

**Figure 5.**
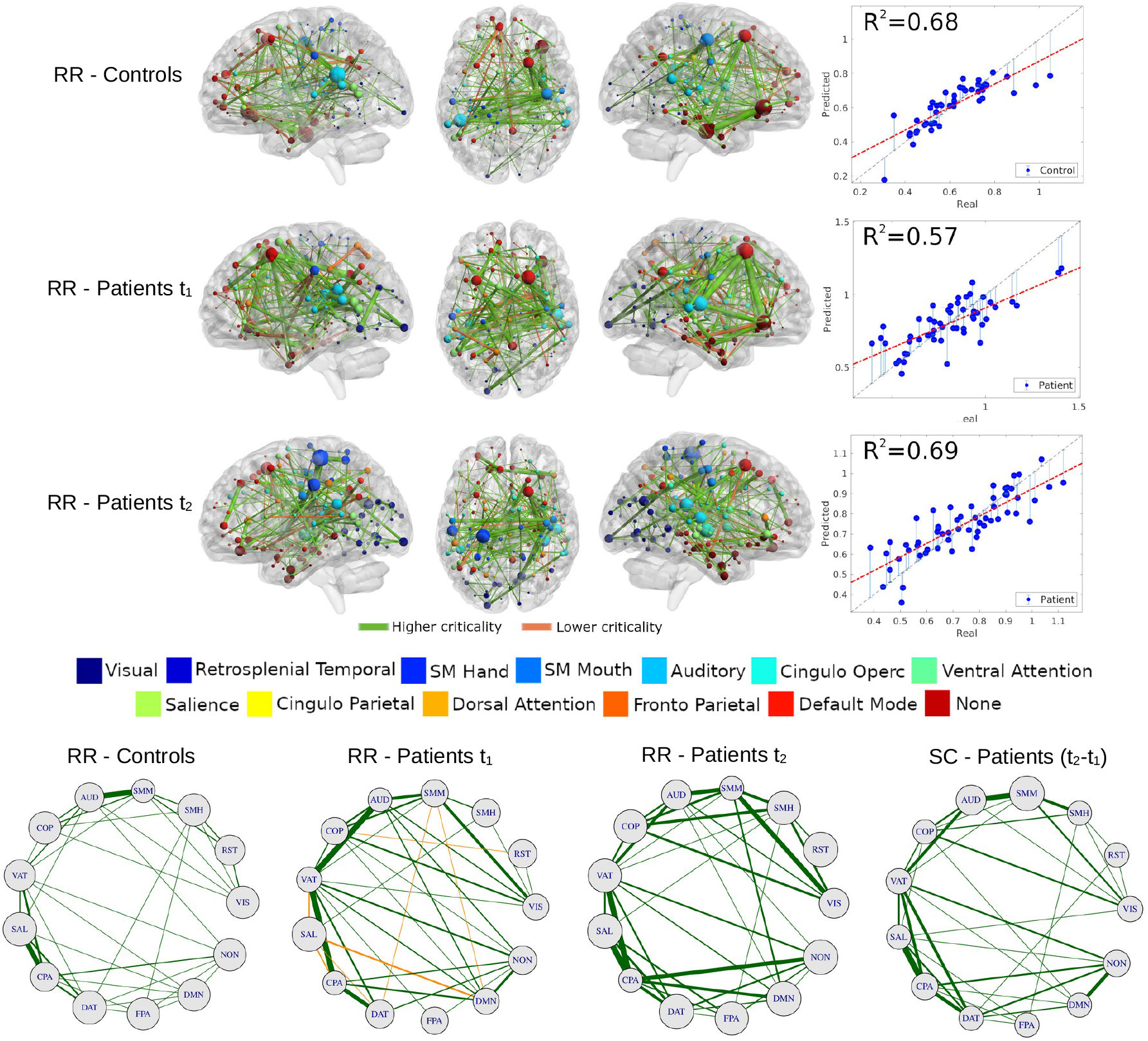
Structural connectivity related to criticality (integral of the second cluster size *I*_2_ = ∫*S*_2_*dT*). Top: map of predictive connections of *W* (see (14)). Structural edges that predict higher (green edges) and lower criticality (orange edges) values (top: controls; middle: patients at *t*_1_; bottom: patients at *t*_2_). Only the highest 200 connections are shown for ease interpretation. The size of each ROI, colored by network, corresponds to the number of predictive edges converging on it. The scatter plot shows real vs. predicted criticality values from the Ridge Regression model. The gray dashed line has a slope of one and is a guide to the eye. Circle plots: architecture of averaged network connections across resting state networks (⟨*W*⟩_*X,Y*_, see main text). The first three plots, from left to right, represents the ridge regression maps (Eq. (14)), while the last map represents the difference of the patients’ structural connectivity matrix at *t*_2_−*t*_1_. The size of each RSNs corresponds to the average connections within networks. Source data are provided as a Source Data file.

The second analysis considers the network topology involving different functional networks (bottom, Fig. 5). Specifically, we analyze the average link between pairs of functional networks, ⟨*W*⟩ _*X,Y*_ = ∑_*i*∈*X*_ ∑ _*j*∈*Y*_ *W*_*i, j*_/*N*_*XY*_, where *X* and *Y* designate ROIs belonging to a given functional network (visual, default mode etc) and *N*_*XY*_ = *N*_*X*_ *N*_*Y*_ is the total number of connections. We use the same sign convention described above for edge colors and show the quartile 50% for ease visualization. The network topology of controls is quite balanced, but few connections are stronger, such as AUD-SMM, SAL-CPA, VAT-CPA, etc. The network topology is dramatically different for patients at *t*_1_. Many pairs of RSNs are associated with criticality loss (orange edges, ⟨*W*⟩ > 0), such as DMN-SAL, DMN-SMM, SAL-VAT, to name a few. There are also more and stronger negative edges between networks than in healthy participants (e.g., VIS-SMM/AUD/COP/VAT). The pattern normalizes on *t*_2_, with more balanced connections compared to *t*_1_, although relatively stronger compared to controls. We used Pearson’s correlation to estimate the similarity between healthy and stroke networks in predicting criticality, as described by the average connectivity across RSNs, ⟨*W*⟩_*XY*_. The correlation between healthy and stroke network at *t*_2_ was moderately high (*ρ*_⟨*W*⟩_ = 0.72), but much smaller at *t*_1_ (*ρ*_⟨*W*⟩_ = 0.36). Both p-values were statistically significant. In summary these findings show that structural connectivity at 12 months post-stroke predictive of criticality becomes more like that of healthy participants.

To examine whether structural connectivity changes predicting criticality truly reflect remodeling of structural connections, we compared, at the level of networks, the ridge regression maps of the edges predicting criticality at 3 or 12 months with the maps of the structural connectivity changes in the same interval The correlation was moderate at 3 months (*ρ* = 0.52), but high at 12 months (*ρ* = 0.80). This result implies that connections predicting criticality values were remodeled over time.

Finally to test if criticality predictive edges were part of the normal functional architecture or reflect random connections, we correlated the number of predictive edges for each node (ROI) with the corresponding node’s strength in the healthy controls’ average functional connectivity. The high correlation seen at both time-points (*ρ* = 0.98 for positive/negative edges vs. node functional connectivity) indicates that the predictive edges are not random, but consistent with the normal variability of the brain functional architecture.

### Relationship between recovery of criticality, functional connectivity, and behavior

Up to this point, our results strongly suggest that stroke recovery induces a normalization of the neural activity patterns that can be quantified by criticality. One important question is whether these dynamical signatures reflect the patients’ recovery as typically seen in behavioral measures^55^ and in the functional connectivity^34^. We used the framework described in^38^ to simulate the functional connectivity from the structural one for each individual patient. Briefly, the time-series of node’s activity, *s*_*i*_(*t*), is convolved with a canonical hemodynamic response function (HRF). We further applied a band-pass filter in the range of 0.01 − 0.1 Hz. Next, we obtain the functional connectivity matrix, FC, through the Pearson correlation, (6), between each pair of ROIs in the network^48^. We use the averaged correlation across ROIs FC to characterize the strength of the functional connections in patients and controls (see (7)). We also compute the entropy, *H*, of the functional matrices following the framework of Saenger et. al.^50^ (see Methods section, (13)). The entropy measures the repertoire diversity and the complexity of the functional connections, and may serve as a biomarker of stroke recovery as well^50^. The behavioral performance of patients and controls was inferred from a neuropsychological battery measuring performance in 8 behavioral domains (motor left */* right, language, verbal and spatial memory, attention visual field, attention average performance, attention shifting)^51^. The Supplementary Note 2 report a full description of the tests used. We used principal component analysis (PCA), a common data reduction strategy that identifies hidden variables or factors, to capture the possible correlation of behavioral scores across participants. In each domain one component explained the majority of variance across participants (57-77% depending on the domain). The component scores were normalized with respect to healthy controls (mean= 0 and SD= 1). This normalization allows to express the patients’ scores in units of standard deviations below average - i.e. *B*(Lang) = −4 is equivalent to language function 4 SD below the control average. Here, to characterize each patient’s overall performance, we use the average component score obtained from averaging the normalized factor scores across domains^51^, ⟨*B*⟩ = ∑_*i*_ *B*_*i*_/8.

The composite behavioral score (*B*) is well behaved as it accurately tracks individual behavioral deficit variability across time points (*R*^2^ = 0.81 and *ρ* = 0.9) (Supplementary Figure 2). To quantify the relationship between dynamical and behavioral deficits we used the integral of the first, *I*_1_ = ∫*S*_1_*dT*, and second, *I*_2_ = ∫*S*_2_*dT*, cluster sizes over the entire range of excitability *T*, respectively. Notably, both variables were strongly correlated with *B* (Fig. 6 A,B,E,F). The correlation is positive for *I*_1_ (*t*_1_ : *ρ* = 0.4; *t*_2_ : *ρ* = 0.5) while it is negative for *I*_2_ (*t*_1_ : *ρ* = −0.25; *t*_2_ : *ρ* = −0.51). This relationship indicates that a larger *S*_1_ cluster and a smaller *S*_2_ cluster are associated with better performance. Note the improvement in correlation of both variables from *t*_1_ to *t*_2_ (compare *t*_1_ Fig. 6 A,E with *t*_2_ B,F). This suggests normalization of the structure-function (behavior) relationship, well captured by the stochastic model over the course of the stroke recovery. Supplementary Table 2 shows the relevant statistical tests corrected for multiple comparisons (FDR). All correlation with behavior were significant except *I*_2_ at *t*_1_ (Fig. 6B). These findings are consistent with the second prediction that variations in neural dynamics are behaviorally relevant.

**Figure 6.**
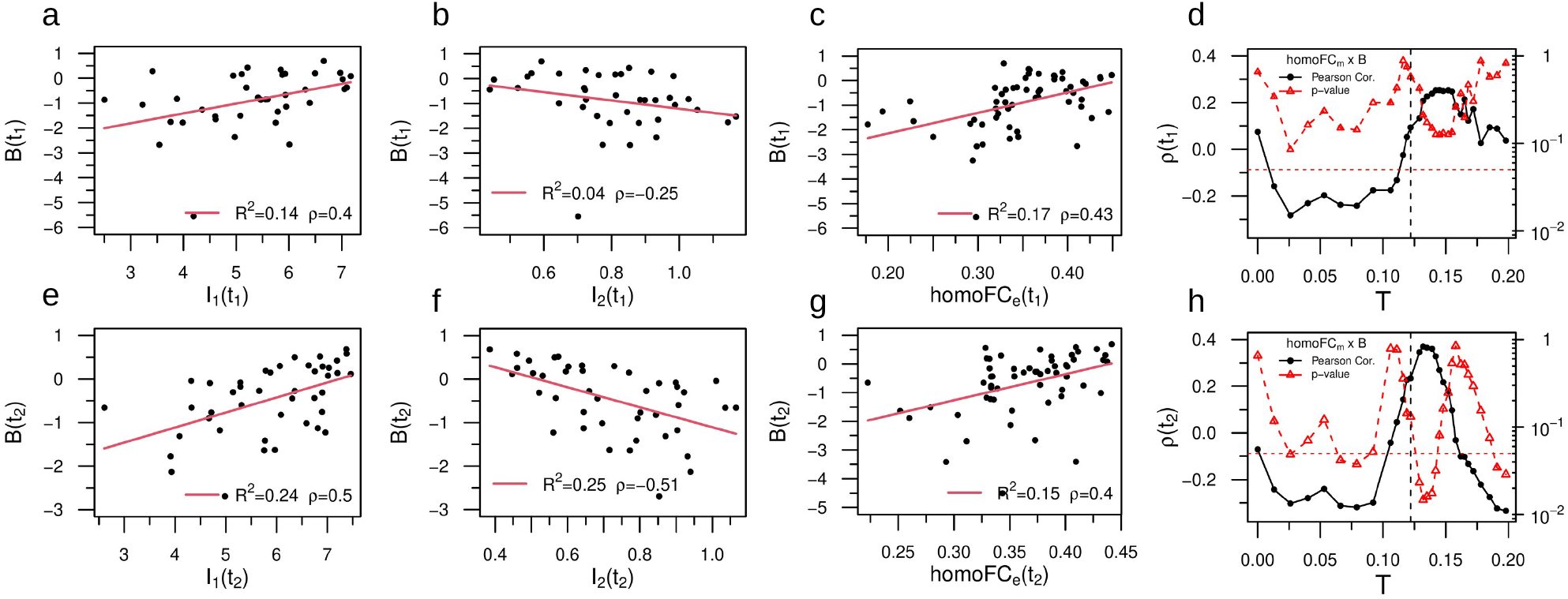
Relationship between recovery of criticality, functional connectivity, and behavior. a,e) Correlation between *I*_1_ and behavior at *t*_1_ and *t*_2_, respectively. The legend shows the (linear) correlation, *ρ* and the *R*^2^. b,f) Same but for *I*_2_. The dynamical variables (*I*_1_, *I*_2_) predicted a significant amount of behavioral variance, higher at *t*_2_. c,g) Correlation between empirical homotopic FC_e_ and behavior at *t*_1_ and *t*_2_, respectively. d,h) Linear correlation between model homo-FC_m_ and behavior for different values of the excitation threshold *T*. Near the critical point (*T*_*c*_, black vertical lines) the stochastic model reproduces the positive correlation with behavior observed empirically (black dots) with a corresponding peak in the statistical significance (red triangles). Note the sharp increase in correlation as *T* nears the critical point and the subsequent fall of as *T* becomes subcritical. We added red horizontal lines at *p* = 0.05 for ease interpretation. All the p-values were Benjamini-Hochberg corrected for multiple comparisons with false discovery rate (FDR) of *α* = 0.05. The p-value reported in panel b did not remain significant after correcting. However, note the abnormal behavior of a single individual close to *B*(*t*_1_) ∼ −6. Removing this outlier from the analysis we obtain *ρ* = −0.42 with p-value < 0.05. See Supplementary Table 2 for accessing the sample size and the significance for all the tests. The (two-tailed) p-values reported in panels (d,h) were not corrected for multiple comparisons. Source data are provided as a Source Data file.

Variations in neural dynamics estimated from the structural model may be reflected in the patterns of functional connectivity (FC). In Fig. 1 we already showed that the model FC derived from applying an hemodynamics response model to the neural dynamics of the whole brain model is weak at sub-critical and super-critical regimes, but strongest at criticality. Here we consider the relationship between model FC and empirically measured FC in the same participants. Studies in stroke have highlighted the behavioral importance of homotopic functional connectivity, i.e. inter-hemispheric connections between symmetrical regions belonging to the same network^37,66,67^. Hence, we examined the relationships between homotopic FC, both empirical - measured directly - and model, with behavioral performance. As a control we used the average FC mediated across all regions. Empirical homotopic FC (homo-FC_e_) correlated significantly with behavioral performance across participants at both time points (*t*_1_ : *ρ* = 0.43 and *t*_2_ : *ρ* = 0.40; p-values < 0.05, corrected; Fig. 6 C-G). As predicted homo-FC_*e*_ showed a stronger correlation with behavior than the average FC_e_ (Supplementary Figure 4). Next, we computed the linear correlation between model homotopic FC (homo-FC_m_) and behavior for the whole range of excitation threshold *T* (Fig. 6 D-H). Notably, we observed at both *t*_1_ and *t*_2_ a sharp increase of the correlation (black dots) close to the critical phase transition (dashed line) that corresponded to a global minimum in the p-value (red dots). In other words, the stochastic whole-brain model poised close to the critical point reproduced the correlation with behavior observed empirically. Statistically significant results were obtained at *t*_2_ (*p* < 0.05), not at *t*_1_. Also note the fall-off of the correlation between the model homotopic FC and behavior as the value of *T* moves toward the subcritical regime. These findings confirm that the model can generate functional connectivity patterns that are like those empirically measured, and that these model FC patterns also correlate with behavioral performance.

To complete the analysis the next step is to compare model neural dynamics with empirical functional connectivity, a well-studied biomarker of stroke and behavior relationships^34,35^. Based on previous studies we employed either the average homo-FC (across all pairs of homotopic regions) or the average FC across all ROIs. We found that variations of the model’s neural activity, as described by *I*_1_ and *I*_2_, were significantly correlated with the empirical average homo-FC_e_ (Fig. 7 A-D). The correlation was positive for *I*_1_ (*t*_1_ : *ρ* = 0.43; *t*_2_ : *ρ* = 0.46; p-values < 0.05, corrected), i.e, large values of *S*_1_ - hence more integrated networks, correlated with the average homotopic connectivity at both time points. On the other side, the correlation was negative for *I*_2_ (*t*_1_ : *ρ* = −0.27; *t*_2_ : *ρ* = −0.44; only the latter p-value remained significant). Interestingly, the relationship with the average empirical FC (FC_e_) was in the same direction, but weaker in correlation strength (Supplementary Table 2): *I*_1_ (*t*_1_ : *ρ* = 0.23; *t*_2_ : *ρ* = 0.29) and *I*_2_ (*t*_1_ : *ρ* = −0.11; *t*_2_ : *ρ* = −0.37). In contrast, we did not find any overall significant correlation between average homo-FC_m_ and average homo-FC_e_ at the critical point *T*_*c*_ (Supplementary Figure 17). Furthermore, the relationship between FC_m_ and FC_e_ not only was maximal at the critical point, but it reached statistical significance at criticality at *t*_2_ (*p* < 0.05), not at *t*_1_ (Supplementary Figure 4). These findings therefore suggest that both model and empirical FC normalize and become more similar consistently with a normalization of structure-function relationships. These results are very encouraging since we did not use any optimization of the model inputs to reproduce the empirical FC as done for example in^49^.

**Figure 7.**
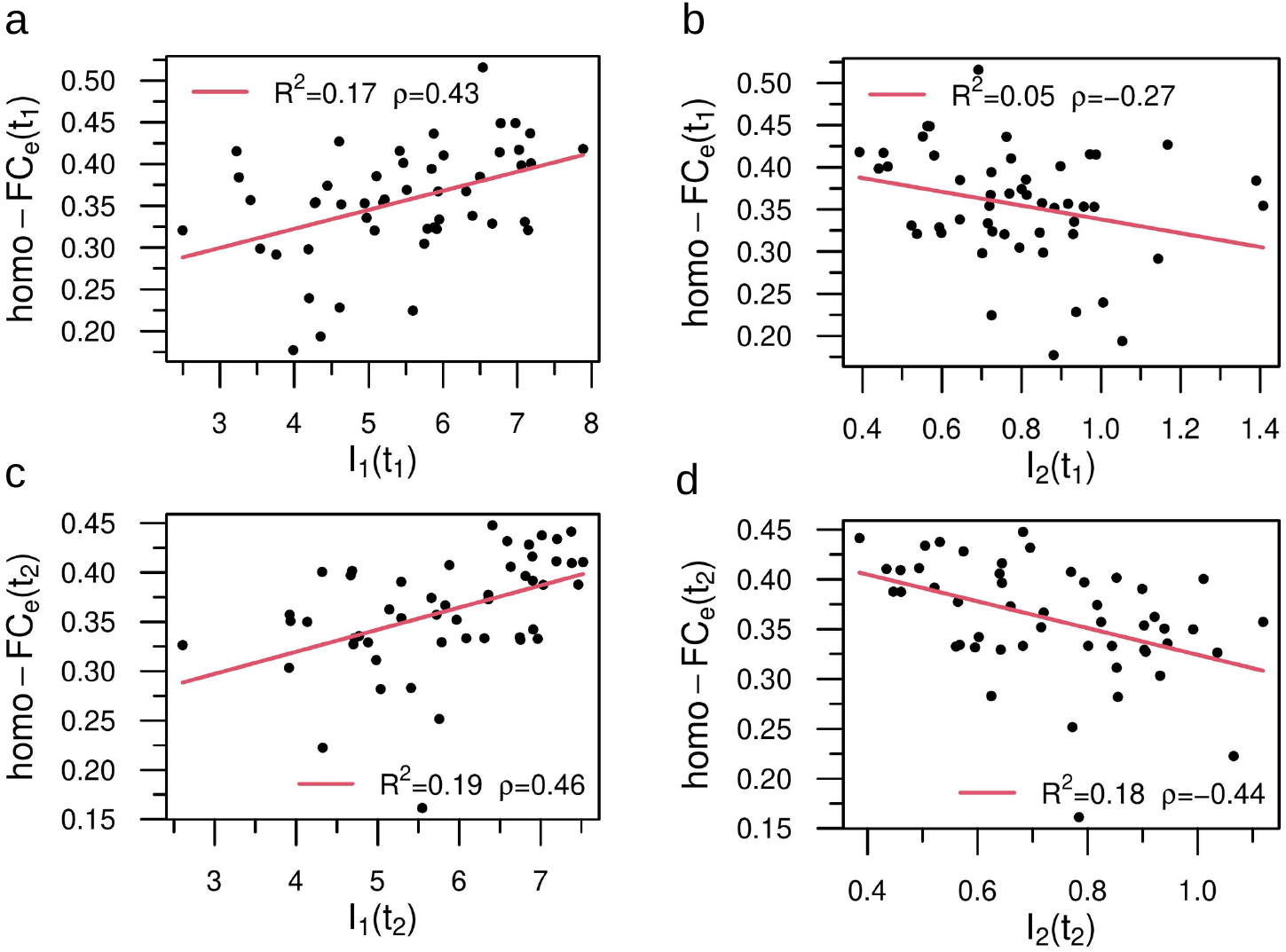
Statistical correlates between model dynamical patterns and empirical homotopic functional connectivity. In the legend we show the (linear) correlation, *ρ* and the *R*^2^. a,c) Relationship between *I*_1_ and empirical homo-FC_e_ at *t*_1_ and *t*_2_, respectively. b,d) The same as before, but for *I*_2_. Both dynamical variables predicted a significant amount of functional variance, but panel b did not remain significant after correcting at level of *α* = 0.05. Sample size (*n* = 50). Source data are provided as a Source Data file.

Overall this section shows that neural dynamics from the model correlate with behavioral performance and that this relationship becomes stronger at 12 months. Moreover, the model can reproduce the patterns of homotopic FC that behave similarly to empirical FC in the correlation with behavior. While it is not surprising given the lack of any optimization that overall empirical FC does not correlate across participants with model FC, the model shows that interestingly empirical and model FC become more similar at criticality, especially when the lesioned model approximates normality in the course of recovery (from 3 to 12 months).

## Discussion

We set out to examine whether criticality is affected by lesions, and whether alterations of criticality are behaviorally relevant. We use lesions as a causal manipulation to test the theory that criticality is a fundamental property of healthy brains that provides optimal functional and behavioral capabilities. Several interesting results are worth of discussion.

First, our stochastic model is personalized since it used as input direct estimates of structural connectivity at the individual level. The model provides measures of activity, functional connectivity, and criticality that tracked individual variability in healthy and stroke participants. Importantly, alterations in stroke patients were evident both at the group and individual level, and easily separated stroke from healthy participants. Second, these criticality alterations normalized over time. This normalization reflect changes of the underlying structural connectivity, with associated changes in the average degree, connectivity disorder, modularity and global efficiency. We describe which connections are most predictive of the final level of criticality, and which predict improvements in criticality. The distribution of predictive connections was not random but matched the normal functional architecture of the healthy brain. Third, we show that alterations of criticality were behaviorally relevant as they correlate with improvements in performance. Finally, we show that the model can reproduce the relationship between functional connections and behavioral deficits/recovery that have been established empirically in many studies^34–36,50,55,66,67^.

### Methodological considerations

The whole-brain mesoscopic model, a variant of the Greenberg-Hastings cellular automata^68^, was proposed by Haimovici et al.^21^. When poised at the critical point, the model captures, at the group level, the emergence of experimental spatiotemporal patterns, the temporal correlation between regions (functional connectivity, FC), the organization of brain wide patterns in so called resting state networks (RSNs), and the scaling law of the correlation length, among others.

We improved the Haimovici model by adding a normalization to each node’s excitatory input, a mechanism of homeostatic plasticity^69–71^. This simple adjustment balanced the macroscopic dynamics increasing the strength of critical transitions. The clusters of activity became more heterogeneous spreading along the whole network and not mainly in the hubs, as in the un-normalized model. In the normalized model, the cluster size distribution in proximity to the critical point follows a truncated power-law with a critical exponent *α* close to the hallmark exponent of avalanches sizes, *α* = 3/2. Finally, the homeostatic normalization mechanism significantly improves the correspondence between simulated and empirical functional networks based on fMRI.

An important feature of the normalized model is that it minimizes the variability of the critical points and neuronal activity patterns among healthy participants. The normalization collapses the model state variables of healthy participants into universal curves, which allows to compare critical points between patients and stroke, and stroke patients at different time points. Whether similar results could be obtained without the inclusion of the homeostatic mechanism is an interesting question that we shall investigate in future work.

Another important innovation is that the input to the model were individually measured structural connectomes, both in healthy participants and stroke patients at two time points. The repeated measures allowed the estimation of the stability of the criticality values that were quite narrow in healthy participants, thus supporting the inference that changes of criticality in stroke were related to the effect of the lesions, and not to the inter-individual variability. The availability of individual structural connectomes is not common, and most whole-brain studies have used population atlases of white matter connections^49,72^. The group of patients studied here is partially in overlap with Lin et al.^73^ who, however, only studied a subgroup of patients with motor deficits. Also Lin et al. used fractional anisotropy, a measure of white matter microstructure in an anatomically defined ROI. Here, for the first time, we use direct tractography creating a structural connectome of each control and stroke patient. This approach is also different from the recent Salvalaggio et al^65^ in which we used an indirect approach to estimate structural disconnections by embedding the lesion in a normative atlas of white matter connections. Another innovative methodological aspect of this work is that we use the structural connectomes without additional optimization - fMRI connectivity is often used to enhance the accuracy of structural connectivity due to its low sensitivity or incomplete coverage^49,50^.

However, the model can be certainly improved. The DWI data were not state-of-the-art. The sequence was 10-year old with 60 directions and a single b-value of 1000 *s*/*mm*^2^. This group of healthy and stroke participants began enrollment in a prospective stroke study at Washington University in 2010 with completion in 2015 (WU Stroke cohort I). The tractography used in this study has been produced following the standards of analyses that lead to the publication of classical atlases of the white matter^74,75^ that match post-mortem dissections^76^ and axonal tracing^77^. Tractography reconstructions were checked by an expert neuroanatomist (MTS). The controls were tested three months apart while the stroke patients were tested at 2 weeks, 3 months and 12 months with the last two time points being the object of this study. We did not acquire diffusion imaging data at 2 weeks given the rapid changes of diffusion signals in the first 1-2 weeks post-stroke. Since most behavioral changes in stroke occur in the first 3 months post-injury, it is likely that the changes in structural connectivity in relation to criticality were actually underestimated. In healthy participants we did not expect significant changes three months apart in their structural connections based on the literature^78,79^. This was empirically confirmed in our healthy control analyses. We have completed the acquisition of a second cohort (WU Stroke cohort II 2016-2020) from which we will have access to multi-shell, multi-directional and multi-weighted diffusion weighted images^80^ at 2 weeks, 3 and 12 months. This will allow us to track structural connectivity changes from 2 weeks to 3 months when the rate of behavioral recovery is the strongest.

We simulated fMRI functional connectivity by augmenting the stochastic whole brain personalized model with a standard hemodynamic model. We used the average (across ROIs) and inter-hemispheric homotopic functional connectivity (homo-FC) and entropy (*H*, which measures the functional weight diversity) to characterize stroke-related changes. The model reproduced changes of functional connectivity observed empirically in stroke, such as a decrease of inter-hemispheric FC^35–37,66,67^ and entropy^50^, subsequent normalization^34^, and correlation with behavioral performance^35^. However, the model’s fit with the empirically measured FC was low (see Supplementary Figure 4 A,C), but approached significance at *t*_2_. We elected not to optimize the input through functional connectivity because it would have hidden the role of structural connectivity in supporting critical phase transitions in stroke patients and longitudinal changes. Fitting the model with free parameters has its own issues including variable parameter sensitivities^81^, identifiability problem^82^ and overfitting issues^83^. More importantly, this work aims at unveiling robust and universal features of brain criticality in relation to the anatomical brain connectivity structure and focal lesions, and therefore it is crucial that the model dynamics has the smallest possible degrees of freedom^84^.

The whole-brain modeling is invariably based on arbitrary choices of the density of structural connectivity and the brain parcellation as well^85^. The latter has been often employed without scrutiny to accommodate the hypotheses of the study in question (e.g., functional versus structural); while the former is subject of intense research and ongoing debates in the literature^86^. A recent study by Jung et al.^85^ shed light on the effect of these two variables on the performance of a whole-brain model represented by a system of interacting oscillators. They reported that the parcellation with different atlases showed similar changes of the architecture of the structural networks, but distinct trends of the goodness-of-fit of the model to the empirical data across tractography densities (i.e, number of streamlines). On the one hand, high densities are desirable to guarantee the reproducibility of the graph-theoretical properties of the structural network; on the other hand, high densities are not always the best condition for whole-brain modeling. These nonlinear effects reflect the inter-individual variability of the brain topologies, thus making it it difficult to have a closed set of tractography parameters that reliably capture the brain architecture. One possible middle ground approach would be based on personalized data processing and modeling^85^.

Along these lines, in the supplementary information (Supplementary Figures 5-9 and Supplementary Note 5) we investigate the sensitivity of the results with respect to the brain parcellation. We test the robustness of our results by replicating our analyses in a uniform parcellation^87^. Specifically, we characterized each brain voxel uniquely based on its anatomical location, in terms of *x, y*, and *z* coordinates. The matrix of anatomical coordinates of the brain was then fed into the k-means clustering in python. We asked the algorithm to parcellate the brain in 400 clusters, to enable the comparison with the Gordon parcellation. In this uniform parcellation all ROIs have equal size, but their functional or anatomical macroscopic relationship have been disorganized, i.e., neighboring voxels can be randomly assigned to other neighboring voxels that can be either correlated or not. However, replication indicates that global neural dynamics parameters like *S*_1_ and *S*_2_ seem to be more dependent on maintenance of local connectivity rather than large scale functional systems.

### Stroke lesions cause changes in activity, entropy, and criticality

Whole-brain models of healthy controls showed stable patterns of neural activity, both across time-points and individuals. It is important to understand the model dynamics in healthy participants before considering changes in stroke. For low thresholds of activation, the system is super-critical with high levels of activity, low entropy, low levels of functional connectivity, and a single giant first cluster (*S*_1_). This is akin to a brain in status epilepticus with very high level of activity but low entropy, hence no efficient processing of information and lack of consciousness. For very high thresholds of activation, the system is sub-critical with low levels of activity, low functional connectivity, and entropy. Activity is mostly local with small clusters (*S*_1_). For intermediate thresholds, the neural patterns followed the expected behavior, with a phase transition peaking around the maximum of *S*_2_ (Fig. 3). In contrast, simulations of the patients’ brains at three months post-stroke show striking attenuation in the signatures of brain criticality for many individuals, as revealed by the patients *S*_2_ monotonic behavior^62^ (see individual profile in the Supplementary Figures 10-16). The curves of overall activity (*A*), variability of activity (*σ*_*A*_), clusters sizes (*S*_1_ and *S*_2_), functional connectivity (FC), and entropy (*H*) are significantly decreased. The first cluster is significantly decreased in size, while the second cluster is significantly larger as compared to controls at multiple thresholds of activation. All these patterns are consistent with lack of criticality^62^. Crucially, the same criticality signatures reveal the recovery at one-year post-stroke for many individuals and on average for the group. The neural patterns at *t*_2_ approach the corresponding controls’ average (see probability distributions of Euclidean distances Fig. 3 and Supplementary Figure 3). Longitudinal changes of the neural variables from *t*_1_ to *t*_2_ were confirmed for *σ*_*A*_, *S*_1_ and *I*_1_.

### Correlation between criticality and behavior

The role of criticality in behavior has been discussed in prior studies^88,89^. For instance, Palva et. al.^90^ reported a correlation between scale-free neuronal avalanches and behavioral time-series in MEG/EEG data. The connection between human cognitive performance and criticality has also been investigated. Ezaki et. al provided empirical support that participants with higher-IQ have neural dynamics closer to criticality than participants with lower-IQ^91^.

Our findings show that focal lesions from stroke, a causal manipulation of brain activity, modify criticality in a significant behavioral manner. We used an aggregate measure of behavioral impairment across multiple domains (motor, vision, attention, memory, language) by averaging factor scores of impairments across domains as in^51^. This aggregate index captured global motor and cognitive disability, as we did not have a specific hypothesis about a link between one behavioral domain and criticality. This performance index manifested known properties such as a strong relationship between different time points, with 3-month scores predicting 12-month scores^55^. It also showed a robust relationship with homotopic inter-hemispheric FC as in previous work^34–36^. Importantly, model neural patterns as indexed by the integral of *S*_1_ and *S*_2_ (*I*_1_, *I*_2_) correlated with behavioral performances at both time points (Fig. 6 A,E / B,F). Patients with higher (normal) performance showed larger *S*_1_ and smaller *S*_2_ clusters. Interestingly, the integral of the cluster sizes, *I*_1_ and *I*_2_, proved a better predictor of behavioral performance than variations in model FC_m_ (Supplementary Figure 4 B,D) or homo-FC_m_ (Fig. 6 D,H).

We find a weak relationship between lesion size and loss of criticality. We did not investigate the effect of lesion size on behavioral dysfunction, a topic that we shall investigate in future work. However, we did not expect a significant influence of lesion size on behavioral deficits and functional/structural connectivity abnormalities post-stroke based on the literature^35,36,51^.

The use of the integrated variables (*I*_1_, *I*_2_) instead of the original ones (*S*_1_, *S*_2_) is motivated by the relevance of the former for clinical applications. First, these integrated variables are highly correlated with the corresponding original variables (*S*_1_, *S*_2_), however, they are more stable. Second, the aberrant behavior of the patients’ neural dynamics is more easily quantified when considering the profile of the curves for all excitation regimes, and not just the value of the cluster’s sizes at the critical point; this latter is more subject to fluctuations from different sources, such as noise in data acquisition, the neural dynamics itself etc. The integrated variables capture relevant biological information as they consider the neural dynamics response to different excitation regimes; this makes it possible to access the response of a brain (from its connectivity matrix) to intrinsic changes in (self) regulatory mechanisms (caused by stroke, for example) that are responsible for the degree of activation and inhibitions. In other words, in general we may have different excitation’s thresholds over the trials (during fMRI acquisition, for instance) and therefore it is a way to conditioning out this degree of freedom that is not accessible from the data.

### Anatomical connections supporting criticality and prediction of recovery

The available theory predicts that criticality deviations are closely related to alterations of the brains network topology^62^. We find statistically significant changes in stroke patients’ average degree, connectivity disorder, modularity and global efficiency as compared to healthy controls (Supplementary Table 3). All these measures exhibited a normalization pattern, with associated probability distributions shifting toward the healthy control distributions (Fig. 4 A-B). Patients at *t*_1_ showed significantly lower average degree, connectivity disorder and global efficiency, but significantly higher modularity. These results are in line with a recent study^92^ performed in a smaller stroke cohort (*n* = 17; patients with more than 6 months since stroke onset) where the authors reported significantly lower global efficiency with significantly higher values of global clustering and modularity as compared to healthy controls. The evolution of these measures over time points was not investigated. In this study we find notable recovery of these global graph measures across time points. Indeed, patients at *t*_2_ tend to normalize the integration-segregation balance exhibiting statistically significant increase in average degree, connectivity disorder, global efficiency and a less robust decrease in modularity. In the Supplementary Information we provide an additional compelling evidence of the intricate relationship between brain criticality, the underlying network topology and behavior (see Supplementary Table 4 and Supplementary Note 3).

Few studies have addressed the relationship between the structural and functional network topology in recovering stroke patients. This same cohort was investigated by Siegel et al.^35^ who analyzed the functional network organization based on fMRI. It is worth noting that the functional modularity presents the opposite trend as reported here; modularity increases over time-points towards the controls; this same study did not find significant differences in global efficiency between controls and patients or over recovery. However, an important difference is that Siegel et al measured modularity in a priori selected functional networks while we measured modularity using a data driven Louvain community detection to optimize the modular organization of the structural network. There are many potential differences that can explain such opposite trend. Functional networks are bilaterally organized, and the breakdown of inter-hemispheric connectivity post-stroke decreases their modularity^35,36^. Moreover, some networks become abnormally linked with other networks, which further decreases network modularity, proportionally to the decrement of inter-hemispheric interaction. In contrast, in a data-driven approach communities will be detected in each hemisphere, early on post-stroke, but their number will decrease in the course of recovery as inter-hemisphere interactions improve.

A notable result is the remodeling of the structural connectivity in the white matter supporting the normalization of criticality from 3 to 12 months post-stroke. Indeed, we find that specific connections in the brain predicted with high accuracy, in a ridge regression model, criticality values (e.g., *I*_2_) at both time points, but with higher accuracy at 12 months. Notably the maps of predictive edges differ from healthy controls at 3 months, normalize at 12 months, but are even more like the map of structural connection changes occurring in the same time interval. In other words, the brain normalizes criticality post-stroke using a different set of connections than a healthy brain. Importantly, these new connections are not randomly located but are part of the normal functional architecture as shown by the strong correlation with the intrinsic pattern of functional connectivity.

The structural connections that predict criticality are best illustrated by the circle plots in Fig. 5. Healthy controls show a balanced set of structural connections predicting criticality (e.g., auditory-ventral attention, ventral attention-cinguloparietal) while others become negatively related to criticality, i.e., their strength predicts low criticality value. Interestingly, some of these latter connections involve the default mode network and other networks (motor, salience). It is speculative to think that this effect may be related functionally to the loss of segregation between regions of the default mode and sensory-motor-attention networks, previously reported^35–37^. At 12 months many of the edges that were negatively predicting criticality (including DMN edges) become positively correlated to criticality. In a recent review, Gollo et. al.^25^ hypothesized that hub regions within the DMN represent a structural signature of near-critical dynamics. Our findings provide some support for this idea. Edges to/from DMN regions as well as networks sub-serving visual, attention and executive control (cingulo-opercular and dorsal attention networks) predicted higher criticality values at *t*_2_.

An increase in critical signatures from *t*_1_ to *t*_2_ must correspond to the recovery of structural (anatomical) connections, which is captured by diffusion-weighted imaging (DWI) and tractography (Supplementary Table 1). Changes in DWI and tractography may reflect a number of different homeostatic plasticity mechanisms, including structural plasticity in gray and white matter tracts, recovery of neural cells, remyelination, and rewiring. Whether long-range anatomical connectional changes support the recovery of function in stroke is not a well-explored issue. Longitudinal changes in micro- and macro-scale structural anatomy and physiology following experimentally induced strokes have been tracked in animals, mostly in the perilesional area^94^. However, there are also observations of long-range plasticity^95–97^. In humans, structural plasticity can be measured at the macro-scale level with diffusion MRI^98^. There is now convincing evidence in both humans and animals that learning through activity-dependent plasticity can modify white matter in healthy adults^99,100^, and possibly in stroke patients^101–103^.

In summary, our theoretical framework to model individual brain dynamics based on real structural connectivity networks suggests that patients affected by stroke present decreased levels of neural activity, decreased entropy, and decreased strength of the functional connections. All these factors contribute to an overall loss of criticality at three months post-stroke that recovers at twelve months, driven by white matter connections remodeling. Notably, our model contains only three parameters (*r*_1_, *r*_2_, and *T*), all set apriori without any fitting procedures. In conclusion, personalized whole-brain dynamical models poised at criticality track and predict stroke recovery at the level of the single patient, thereby opening promising paths for computational and translational neuroscience.

## Methods

This research complies with all relevant ethical regulations. All studies with human participants were approved by the Washington University Institutional Review Board. Stroke patients and healthy controls provided informed consent according to procedures approved by the Washington University Institutional Review Board.

### Stroke dataset

All data came from a large prospective longitudinal stroke study described in previous publications^35,51,55^. We provide here a brief description of the dataset and refer the reader to those articles for a more comprehensive description.

Clinical sample: The dataset includes 132 stroke patients (mean age 54, standard deviation 11, range 19-83; 71 males; 68 left side lesions) at the sub-acute stage (2 weeks post-stroke). The inclusion/exclusion criteria were as follows: first symptomatic stroke, ischemic or hemorrhagic, and clinical evidence of any neurological deficit. We used data from the subset of 103 patients who returned for clinical and imaging assessments at three months post-stroke, as well as the data from the 88 patients who returned for 1 year post-stroke assessment (for details see Corbetta et al.^51^). The control group, formed by 28 individuals, was matched with the stroke sample for age, gender, and years of education. Data was collected twice in the healthy controls, 3 months apart.

The neuropsychological battery included 44 behavioral scores across five behavioral domains: language, memory, motor, attention, and visual function. These domains were chosen to represent a wide range of the most commonly identified deficits in people after a stroke.

### MRI Acquisition

Patients were studied 2 weeks (mean = 13.4 d, SD = 4.8 d), 3 months (mean = 112.5 d, SD = 18.4 d), and 1 year (mean = 393.5 d, SD = 55.1 d) post-stroke. Diffusion data were obtained only at 3 months and 1 year. Controls were studied twice with an interval of 3 months. All imaging was performed using a Siemens 3T Tim-Trio scanner at WUSM and the standard 12-channel head coil. The MRI protocol included structural, functional, pulsed arterial spin labeling (PASL) and diffusion tensor scans. Structural scans included: (i) a sagittal T1-weighted MPRAGE (TR=1,950 ms, TE=2.26 ms, flip angle=90°, voxel size= 1.0 × 1.0 × 1.0 mm); (ii) a transverse T2-weighted turbo spin echo (TR = 2,500 ms, TE = 435 ms, voxel size = 1.0 × 1.0 × 1.0 mm); and (iii) sagittal fluid attenuated inversion recovery (FLAIR) (TR = 7,500 ms, TE = 326 ms, voxel size = 1.5 × 1.5 × 1.5 mm). PASL acquisition parameters were: TR = 2,600 ms, TE = 13 ms, flip angle = 90°, bandwidth 2.232 kHz/Px, and FoV 220 mm; 120 volumes were acquired (322 s total), each containing 15 slices with slice thickness 6- and 23.7-mm gap. Resting state functional scans were acquired with a gradient echo EPI sequence (TR = 2,000 ms, TE = 27 ms, 32 contiguous 4-mm slices, 4 × 4 mm in-plane resolution) during which participants were instructed to fixate on a small cross in a low luminance environment. Six to eight resting state fMRI runs, each including 128 volumes (30 min total), were acquired. fMRI Data Preprocessing of fMRI data included: (i) compensation for asynchronous slice acquisition using sinc interpolation; (ii) elimination of odd/even slice intensity differences resulting from interleaved acquisition; (iii) whole brain intensity normalization to achieve a mode value of 1,000; (iv) removal of distortion using synthetic field map estimation and spatial realignment within and across fMRI runs; and (v) resampling to 3-mm cubic voxels in atlas space including realignment and atlas transformation in one resampling step. Cross-modal (e.g., T2 weighted to T1 weighted) image registration was accomplished by aligning image gradients. Cross-model image registration in patients was checked by comparing the optimized voxel similarity measure to the 97.5 percentile obtained in the control group. In some cases, structural images were substituted across sessions to improve the quality of registration.

Diffusion weighted imaging (DWI) included a total of 64 near-axial slices. We used a fully optimized acquisition sequence for tractography that provided isotropic (2 × 2 × 2 mm) resolution and coverage of the whole head with a posterior-anterior phase of acquisition. We set the echo time (TE) and the repetition time (TR) to 9.2 milliseconds and 9200 milliseconds, respectively. At each slice location, 4 images were acquired with no diffusion gradient applied. Additionally, 60 diffusion-weighted images were acquired, in which gradient directions were uniformly distributed on the hemisphere with electrostatic repulsion. The diffusion weighting was equal to a b-value of 1000 sec mm2. In order to optimize the contrast of acquisition, this sequence was repeated twice.

### MRI and Lesion Analysis

Individual T1 MRI images were registered to the Montreal Neurological Institute brain using FSL (FMRIB Software Library) FNIRT (FMRIB nonlinear imaging registration tool). Lesions were manually segmented on individual structural MRI images (T1-weighted MPRAGE, T2-weighted spin echo images, and FLAIR images obtained 1–3 wk post-stroke) using the Analyze biomedical imaging software system (www.mayoclinic.org). Two board-certified neurologists (M.C. and Alexandre Carter) reviewed all segmentations. Special attention was given to distinguish lesion from cerebral spinal fluid (CSF), hemorrhage from surrounding edema, and to identify the degree of periventricular white matter damage present. In hemorrhagic strokes, edema was included in the lesion. A neurologist (M.C.) reviewed all segmentations a second time, paying special attention to the borders of the lesions and degree of white matter disease. The staff that was involved in segmenting or in reviewing the lesions was blind to the individual behavioral data. Atlas-registered segmented lesions ranged from 0.02 to 82.97 cm3 with a mean of 10.15 cm3 (SD = 13.94 cm3). Lesions were summed to display the number of patients with structural damage for each voxel.

### Functional Connectivity (FC) Processing

FC processing followed previous work from the laboratory (see^34^), with the addition of surface projection and processing steps developed by the Human Connectome Project. First, data were passed through several additional preprocessing steps: (i) regressors were computed based on Freesurfer segmentation; (ii) removal by regression of the following sources of spurious variance: (a) six parameters obtained by rigid body correction of head motion, (b) the signal averaged over the whole brain, signal from ventricles and CSF, and (d) signal from white matter; (ii) temporal filtering retaining frequencies in the 0.009–0.08Hz band; and (iii) frame censoring. The first four frames of each BOLD run were excluded. Frame censoring was computed using framewise displacement with a threshold of 0.5 mm. This frame-censoring criterion was uniformly applied to all R-fMRI data (patients and controls) before functional connectivity computations. Participants with less than 120 usable BOLD frames were excluded (13 patients, 3 controls).

Surface generation and processing of functional data followed procedures similar to Glasser et al.^105^, with additional consideration for cortical segmentation in stroke patients. First, anatomical surfaces were generated for each participant’s T1 MRI using FreeSurfer automated segmentation^106^. This included brain extraction, segmentation, generation of white matter and pial surface, inflation of the surfaces to a sphere, and surface shape-based spherical registration to the participants (native) surface to the fs average surface. Segmentations were manually checked for accuracy. For patients in whom the stroke disrupted automated segmentation, or registration, values within lesioned voxels were filled with normal atlas values before segmentation, and then masked immediately after (seven patients). The left and right hemispheres were then resampled to 164,000 vertices and registered to each other, and finally downsampled to 10,242 vertices each for projection of functional data. Following preprocessing of BOLD data, volumes were sampled to each participant’s individual surface (between white matter and pial surface) using a ribbon-constrained sampling available in Connectome Workbench. Voxels with a high coefficient of variation (0.5 SDs above the mean coefficient of variation of all voxels in a 5-mm sigma Gaussian neighborhood) were excluded from volume to surface mapping^105^. Time courses were then smoothed along the 10,242 vertex surface using a 6-mm FWHM Gaussian kernel. Finally, time courses of all vertices within a parcel are averaged to make a parcelwise time series. We used a cortical surface parcellation generated by Gordon et al.^59^. The parcellation is based on R-fMRI boundary mapping and achieves full cortical coverage and optimal region homogeneity. The parcellation includes 324 regions of interest (159 left hemisphere, 165 right hemisphere). The original parcellation includes 333 regions, and all regions less than 20 vertices (approximately 50 mm2) were excluded. Notably, the parcellation was generated on young adults age 18–33 and is applied here to adults age 21–83.

### Diffusion weighted imaging (DWI) processing

For each slice, diffusion-weighted data were simultaneously registered and corrected for participant motion and geometrical distortion adjusting the diffusion directions accordingly^107^ (ExploreDTI http://www.exploredti.com). Spherical deconvolution was chosen to estimate multiple orientations in voxels containing different populations of crossing fibres^108–110^. The damped version of the Richardson-Lucy algorithm for spherical deconvolution^57^ was calculated using Startrack (https://www.mr-startrack.com).

Algorithm parameters were chosen as previously described^56^. A fixed fibre response corresponding to a shape factor of *α* = 1.5 × 10^−3^*mm*^2^*/s* was chosen^56^. Fibre orientation estimates were obtained by selecting the orientation corresponding to the peaks (local maxima) of the fibre orientation distribution (FOD) profiles. To exclude spurious local maxima, we applied both an absolute and a relative threshold on the FOD amplitude. A first absolute threshold was used to exclude intrinsically small local maxima due to noise or isotropic tissue. This threshold was set to 3 times the mean amplitude of a spherical FOD obtained from a grey matter isotropic voxel (and therefore also higher than an isotropic voxel in the cerebrospinal fluid). A second relative threshold of 10% of the maximum amplitude of the FOD was applied to remove the remaining local maxima with values higher than the absolute threshold^111^.

Whole-brain tractography was performed selecting every brain voxel with at least one fibre orientation as a seed voxel. From these voxels, and for each fibre orientation, streamlines were propagated using Euler integration with a step size of 1 mm (as described in^56^). When entering a region with crossing white matter bundles, the algorithm followed the orientation vector of least curvature (as described in Schmahmann and Pandya^112^). Streamlines were halted when a voxel without fibre orientation was reached or when the curvature between two steps exceeded a threshold of 60°.

Normalization to the MNI152 space was performed after reconstructing the streamline in the native space of the patients^113^. We co-registered the structural connectome data to the standard MNI 2 mm space using the following steps: first, whole-brain streamline tractography was converted into streamline density volumes where the intensities corresponded to the number of streamlines crossing each voxel. Second, individual streamline density volumes were registered to the streamline density template in the MNI152 space template derived from^60^ masking for the lesion^114,115^ and the same transformation was applied to the individual whole-brain streamline tractography using the trackmath tool distributed with the software package Tract Querier^116^. Hence uniform deformation was applied to the whole brain and did not produce distortion that mostly occur when applying T1w normalisation to tractography. Further quality of the streamline normalisation was visually inspected by an anatomist (MTS).

Dissections were performed using trackvis^117^ (http://trackvis.org). Regions of interest were derived from^59^ and arranged 2 by 2 in order to select streamlines and build a connectivity matrix for each patient. We considered the number of streamlines existing between two regions as a surrogate of the strength of the connection. Although the number of streamlines is not precise enough for an accurate estimate of fibre strength^118^, it is acceptable in the context of brain disconnection after a stroke^119,120^.

Our tractography approach did not set constraints on connectivity density. The individual connectivity matrices included very weak connections and this prevented the model from showing criticality. Indeed, as demonstrated by Zarepour et al.^62^, the model shows a first-order phase transition at high densities, no phase transition at low density, and it is critical for intermediate values. We therefore removed the weakest connections (≤ 3 streamlines) from each individual matrix (both for patients and controls), which was sufficient to reproduce the expected critical behavior.

### Characterization of simulated brain activity

We have considered the following standard quantities to characterize the simulated brain activity:

- the mean network activity,

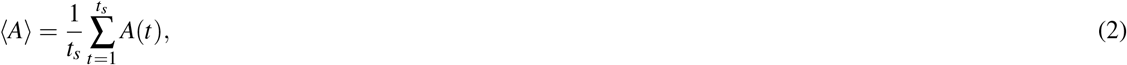

where 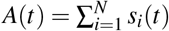 is the instantaneous activity and *t*_*s*_ is the simulated total time;
- the standard deviation of *A*(*t*),

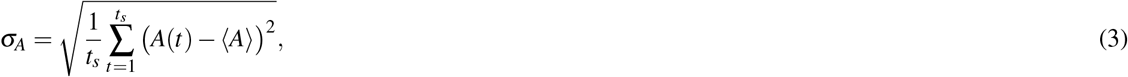
- the sizes of the averaged clusters, the largest ⟨*S*_1_⟩ and the second largest ⟨*S*_2_⟩. Clusters were defined as ensembles of nodes that are structurally connected to each other and simultaneously active. We use numerical derivative (forward finite difference method) to determine the monotonic behavior of *S*_2_. In theory, the monotonic behavior occurs when the sign of the derivative remains constant. We relaxed this criterion due to the noisy behavior of the derivative / model data (mainly for small values of *T*). Therefore, for a few cases where the change of sign happened before *T* ≲ 0.07 we attribute it as noise or artifact, and classified it as monotonic behavior.

Following our previous work^38^, we set the model parameters to the following values, *r*_1_ = 2*/N* (with *N* = 324), 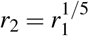, and we vary the activation threshold *T* ∈ [0, 0.2]. We updated the network states, starting from random configurations of *A, I* and *R* states, for a total of *t*_*s*_ = 2000 time-steps. For each value of the threshold *T* we computed the state variables, ⟨*S*_1_⟩, ⟨*S*_2_⟩, ⟨*A*⟩ and *σ*_*A*_. Throughout this study, unless stated otherwise, the final numerical results presented were averages over 10 initial random configurations. For computation of model data, we discarded the initial transient dynamics (first 100 time steps).

### From the model output to BOLD signal

We have employed a standard procedure to transform model output in BOLD functional signals^21,38^. Accordingly, the node’s activity, *s*_*i*_(*t*), is convolved with a canonical double-gamma hemodynamic response function (HRF),

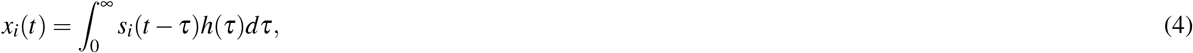

with,

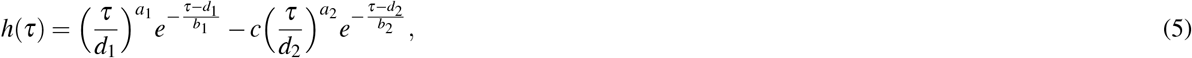

where *x*_*i*_(*t*) is the BOLD signal of the *i*-th node. The free parameters in (5) were fixed according to values found in^121^, i.e., *d*_*i*_ = *a*_*i*_*b*_*i*_, *a*_1_ = 6, *a*_2_ = 12, *b*_*i*_ = 0.9, and *c* = 0.35. Finally, the BOLD time-series, **x**(*t*), were filtered with a zero lag finite impulse response band pass filter in the frequency range of 0.01 − 0.1 *Hz*.

From the generated BOLD signal we can finally extract the following quantities:

- the functional connectivity network (FC). In fact, the FC matrix FC_*i j*_ is defined through Pearson correlation:

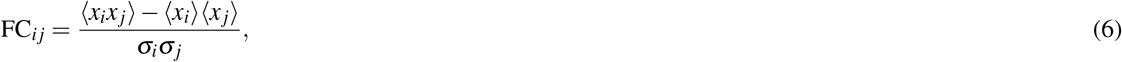

where 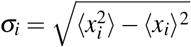 is the standard deviation and ⟨·⟩ is the temporal average of the BOLD time series.
- the average of the functional connectivity:

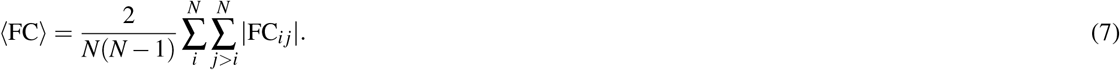

From the above expression we observe that only the upper triangular elements of |FC| are considered in the average.
- the Shannon entropy:

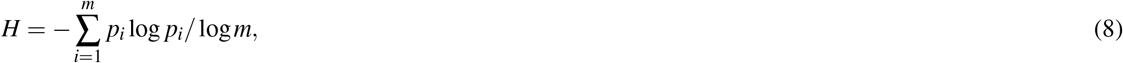

where *m* is the number of bins used to construct the probability distribution function of the upper triangular elements of |FC|. The normalization factor in the denominator, i.e., log *m*, is the entropy of a uniform distribution, and it ensures that *H* is normalized between 0 and 1. The distributions were partitioned with *m* = 20 bins. The higher the diversity of the functional connectivity matrix, the higher the entropy of that functional connectivity matrix.
- finally, we characterize the distance between any given simulated neural variable with the corresponding controls average through the Euclidean distance:

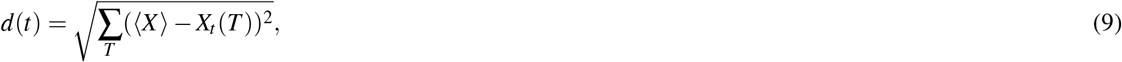

where *X*_*t*_(*T*) is a given neural pattern at time-point *t* and threshold *T*, while ⟨*X*⟩ is the corresponding controls average.

### Network measures of brain connectivity

We have applied the following measures from graph theory to quantify the topology of the corresponding structural brain networks^64^:

- Newman’s modularity was calculated from the binary (undirected) adjacency matrix (*A*_*vω*_):

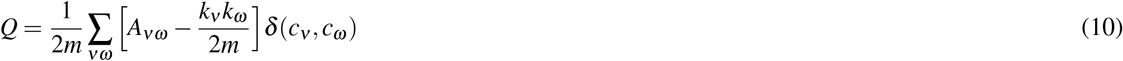

where the network is fully subdivided into a set of nonoverlapping communities such that node *v* belongs to community 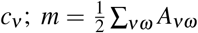 is the number of edges and *k*_*v*_ = ∑_*ω*_ *A*_*vω*_ is the degree of node *v*. We employed the Louvain community detection as implemented in R (igraph package).
- The global efficiency:

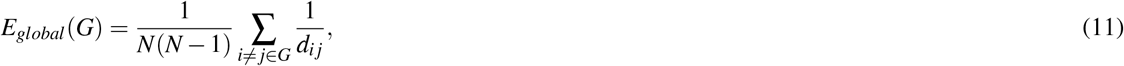

where *d*_*i j*_ is the shortest path length between nodes *i* and *j*. We used the (binary) adjacency matrix to compute the shortest path lengths.
- The average degree:

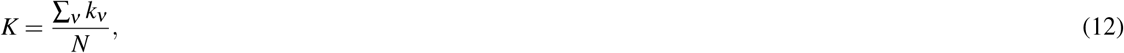

where *N* is the number of nodes and *k*_*v*_ is the degree of node *v* as defined above.
- The connectivity disorder (i.e., structural entropy):

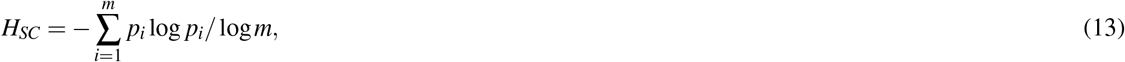

where *m* = 100 is the number of bins used to construct the probability distribution function of all the elements of 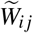.

### Mapping criticality to structural connectivity

The main aim of these analyses was to identify topographical patterns of the structural connectivity matrix (SC) that are related to criticality indexes through multivariate (machine learning) analyses. In our multivariate approach (also see Siegel et al.^35^ and Salvalaggio et al.^65^), features of the individual SC matrices extracted by Principal Component Analysis (PCA) were used as multivariate predictors for a Ridge Regression (RR) model trained to predict patients’ criticality values. RR differs from multiple linear regression because it uses L2-normalization to regularize model coefficients, so that unimportant features are automatically down weighted or eliminated, thereby preventing overfitting and improving generalization on test data^122^. The model weights *W* are computed as:

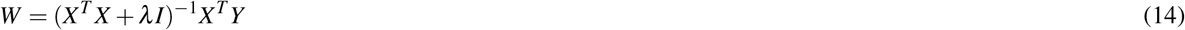

where *X* is the set of predictors and *Y* is the outcome variable. The regularization term provides a constraint on the size of weights and it is controlled by parameter *λ*. A tuning procedure is necessary to find the appropriate value of *λ*. Importantly, this approach also allows to project predictive weights back to brain data in a very simple way^35,123^. Before applying RR, principal component analysis (PCA) was performed on the SC matrix to reduce the input dimensionality. The latter included 52, 326 edges, corresponding to all non-diagonal elements of one half of the symmetric SC matrix of 324 nodes/parcels. Principal Components (PCs) that explained 95% of the variance were retained and used as input for the RR model. All predictors (PC scores) and the outcome variable (criticality value) were z-normalized before applying RR. All RR models were trained and tested using a leave-one-(patient)-out cross validation (LOOCV) loop^124^. In each loop, the regularization coefficient *λ* was optimized by identifying a value between *λ* = 10^−5^ and 10^5^ (logarithmic steps) that minimized leave-one-out prediction error over the training set. Optimal weights were solved across the entire training set using gradient descent to minimize error for the ridge regression equation by varying *λ*. These model weights were then applied to the left-out patient to predict the criticality value. A prediction was generated for all patients in this way. Model accuracy was assessed using the coefficient of determination

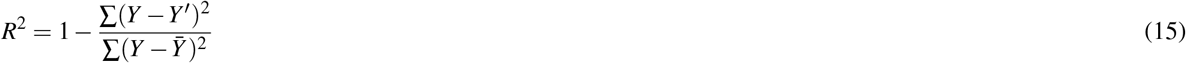

where *Y* are the measured criticality values, *Y*′ are the predicted criticality values and 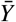 is the mean of predicted criticality indexes. The statistical significance of all LOOCV results was assessed using permutation test. For each regression model, criticality scores were randomly permuted across participants 10000 times, and the entire regression was carried out with each set of randomized labels. A P-value was calculated as the probability of observing the reported *R*^2^ values by chance (number of permutation *R*^2^ > observed *R*^2^)/(number of permutation). Finally, the RR weight matrix was averaged across all LOOCV loops to generate a single set of consensus weights. Statistical reliability of each consensus weight was assessed by comparing its distribution of values (throughout the LOOCV loops) to a null distribution (obtained from the null models generated for permutation testing) using a FDR corrected t-test. The final set of (statistically reliable) consensus weights was back projected to the brain to display a map of the most predictive structural connections.

## Data availability

Data to replicate all the figures and tables are provided as Source data with this paper and are also deposited in Github (https://github.com/CorbettaLab/Rodrigo2022NatComm) and Zenodo^125^ (https://doi.org/10.5281/zenodo.6459955). Individual structural connectivity matrices for controls and patients have been deposited in the Github and Zenodo repositories. Raw neuroimaging and neuropsychological data from^35,51^ are publicly available at cnda.wustl.edu and require controlled access as they contain sensitive patients’ data. The person requesting the data must sign a confidentiality agreement provided by Washington University stipulating that they will make no attempt to identify the patients and to use data only for research purposes. Correspondence and requests for materials should be addressed to R.P.R.

## Code availability

The custom code for criticality modeling is freely available at https://github.com/CorbettaLab/Rodrigo2022NatComm and https://doi.org/10.5281/zenodo.6459955.

## Acknowledgements

We would like to express our gratitude to Professor Sergio Souto Rocha (in memoriam) for his insightful advice and discussions during all stages of this research. R.P.R. was supported by the Research, Innovation and Dissemination Center for Neuro-mathematics (FAPESP Grant No. 2018/08609-8) and by the National Council for Scientific and Technological Development (CNPq Grant No. 201241/2015-3). M.D.F. and M.Z. were supported by the Italian Ministry of Health under Grant Number RF-2013-02359306 to M.Z.

## Author Contributions Statement

R.P.R. and M.C. conceived the study. M.T.S. performed the pre-processing of structural connectivity data. R.P.R. performed the research. M.D.F. and M.Z. performed machine learning and statistical analyses. R.P.R. analyzed the model data and discussed the results with L.K., S.S., M.D.F., M.T.S., M.Z. and M.C. R.P.R. wrote the first draft of the manuscript and all authors reviewed the manuscript.

