## Supplementary Information for "Recovery of neural dynamics criticality in personalized whole brain models of stroke"

[1]Departamento de Física, Centro de Ciências Físicas e Matemáticas, Universidade Federal de Santa Catarina, 88040-900, Florianópolis, SC, Brazil.

[2]Department of Physics, School of Philosophy, Sciences and Letters of Ribeirão Preto, University of São Paulo, Ribeirão Preto, SP, Brazil.

[3]Padova Neuroscience Center, Università di Padova, Padova, Italy.

[4]Laboratory of Neural Computation, Istituto Italiano di Tecnologia, 38068 Rovereto, Italy.

[5]Dipartimento di Fisica e Astronomia, Università di Padova and INFN, via Marzolo 8, I-35131 Padova, Italy.

[6]Fondazione Ospedale San Camillo IRCCS, Venezia, Italy.

[7]Brain Connectivity and Behaviour Laboratory, BCBlab, Sorbonne Universities, Paris France.

[8]Groupe d’Imagerie Neurofonctionnelle, Institut des Maladies Neurodégénératives-UMR 5293, CNRS, CEA University of Bordeaux, Bordeaux, France.

[9]Dipartimento di Psicologia Generale, Università di Padova, Padova, Italy.

[10]Dipartimento di Neuroscienze, Università di Padova, Padova, Italy.

[11]Venetian Institute of Molecular Medicine (VIMM), Fondazione Biomedica, Padova

### Supplementary Figures

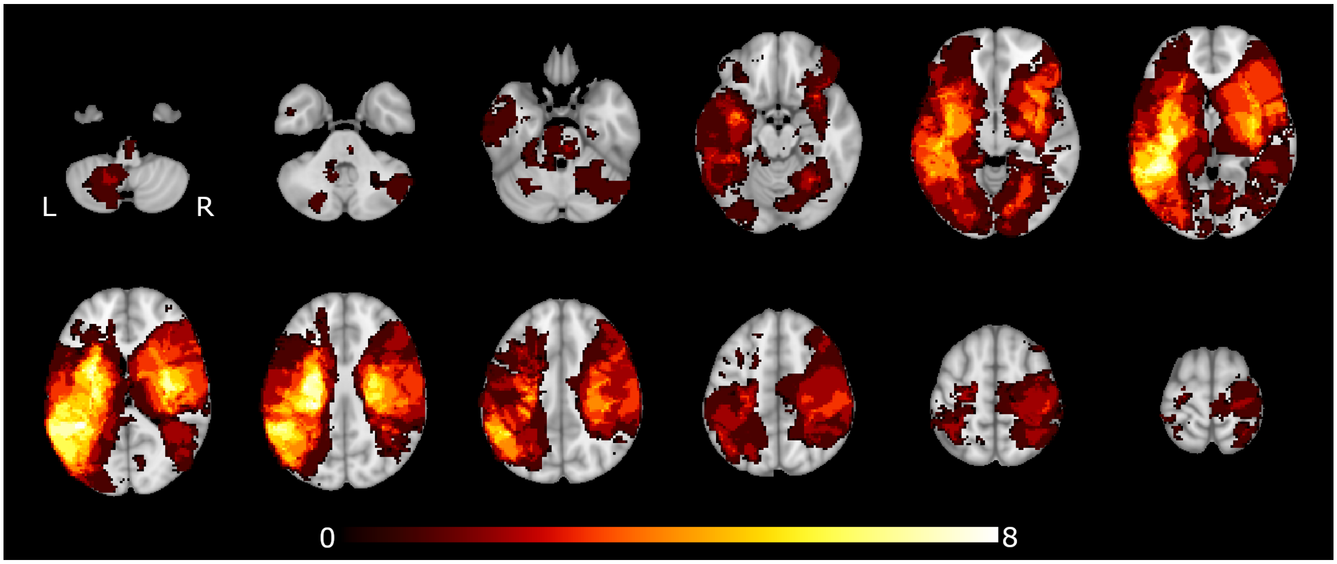


**Supplementary Figure 1** Lesion distribution of the stroke cohort ($n = 56$). Color scale indicates number of patients with a lesion in a given voxel.

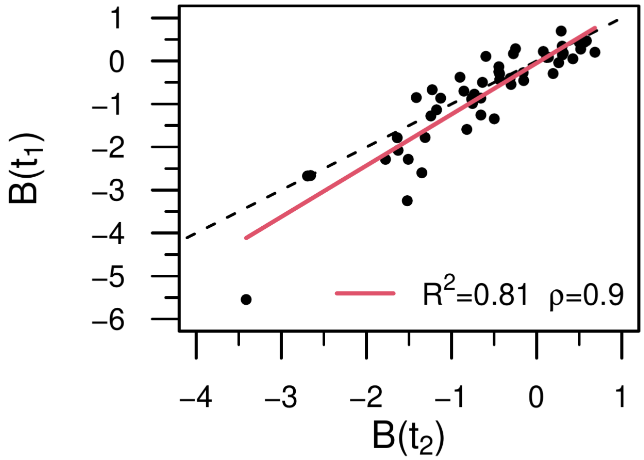


**Supplementary Figure 2** Behavioral deficit variability across time points. Average factor score ($B$) at $t_1$ and $t_2$. In the legend we show the (linear) correlation, $\rho$ and the $R^2$. The gray dashed line has a slope of one and is a guide to the eye. Sample size ($n = 51$).

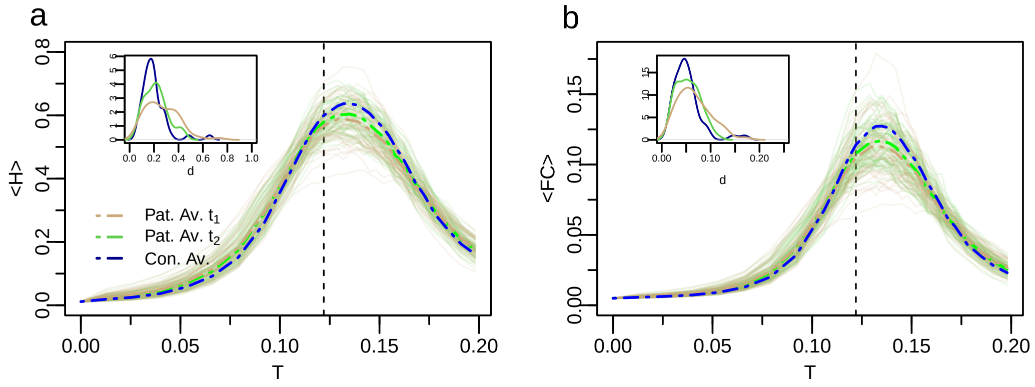


**Supplementary Figure 3** Decreased functional connectivity and entropy in stroke patients. a-b) Group based analysis of neural activity patterns, entropy ($H$) and average functional connectivity (FC), as a function of $T$ for all patients and controls (blue lines). Brown lines represent patients at $t_1$ (3 months post-stroke, $n = 54$), while green lines at $t_2$ (12 months post-stroke, $n = 59$). The thin solid curves represent each individual stroke patient, while the heavy dotted lines represent the group average. Insets: Probability distribution function (pdf) of the Euclidean distances $d$ in individual age-matched-controls (blue) and stroke patients at $t_1$ (brown) and $t_2$ (green). Patients at $t_1$ show greater variability in model neural activity as depicted by the longer pdf's tails. Note the trend toward normalization from $t_1$ to $t_2$.

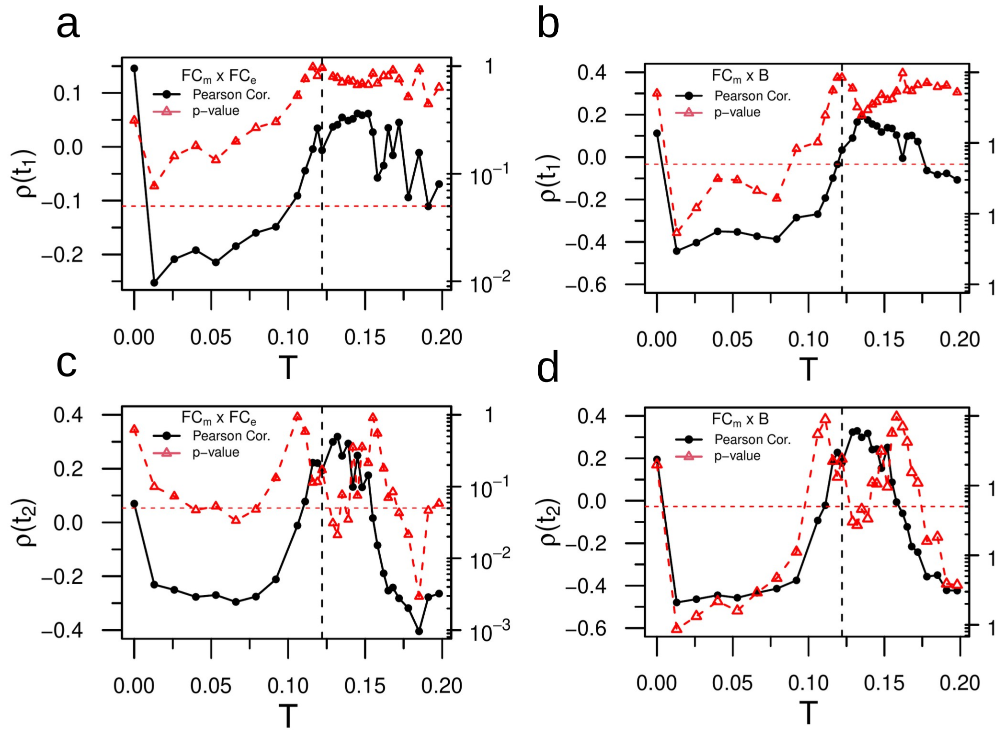


**Supplementary Figure 4** Relationship between functional connectivity and behavior at criticality. a,c) Linear correlation (two-tailed) between model $FC_m$ and empirical $FC_e$ for various values of the excitation threshold $T$ for stroke patients at $t_1$ and $t_2$, respectively. b,d) The same as the previous one, but for $B$. Note the sharp increase in correlation (black dots) close to the critical point and associated behavior of the p-values (red dots). We added red horizontal lines at $p = 0.05$ for ease interpretation. Only the second time-point is significant close to the critical point. The p-values reported in panels a-d were not corrected for multiple comparisons.

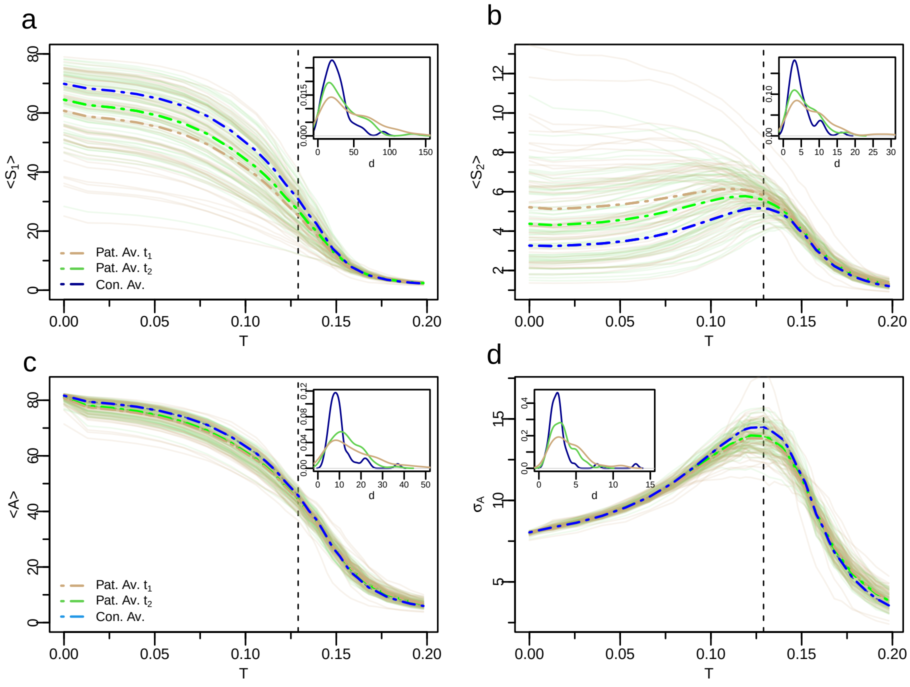


**Supplementary Figure 5** Sensitivity of the results in a uniform parcellation. Variability of the neural dynamics criticality across groups and time points. a,b) Group based analysis of neural activity patterns, $S_1$ and $S_2$, as a function of $T$ for all patients and controls. The brown lines represent patients at $t_1$ (3 months post-stroke), and the green lines at $t_2$ (12 months post-stroke). The thin solid curves indicate each individual stroke patient, while the thick dashed lines represent the group average. c,d) Average activity and standard deviation of activity, respectively. Note improvement of the activity and its variability across time points. Insets: Probability distribution function (pdf) of the Euclidean distances $d$ in individual age-matched-controls and stroke patients at $t_1$ (brown) and $t_2$ (green). The similarity with the Gordon's parcellation (main text, Fig. 3) is quite evident.

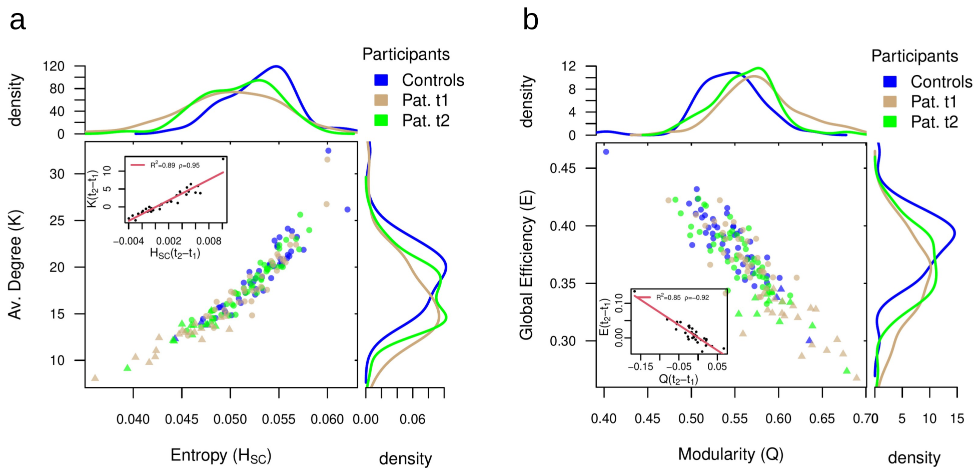


**Supplementary Figure 6** Sensitivity of the results in a uniform parcellation. Alterations in network topology and associated changes in criticality regime. a) Average degree ($K$) versus connectivity disorder (i.e., structural entropy $H_{SC}$) for all patients and controls. Controls are colored in blue. Brown dots represent patients at $t_1$ (3 months post-stroke), while green dots at $t_2$ (12 months post-stroke). Normalized density plots (histograms) are also shown. Triangular dots correspond to patients who have $S_2$ monotonic decay and so lack of criticality. In the inset we show the recovery indexes (see main text) $K(t_2 - t_1)$ versus $H_{SC}(t_2 - t_1)$. In the legend we show the (linear) correlation, $\rho$ and the $R^2$. b) The same as a) but for the global efficiency ($E$) and modularity ($Q$).

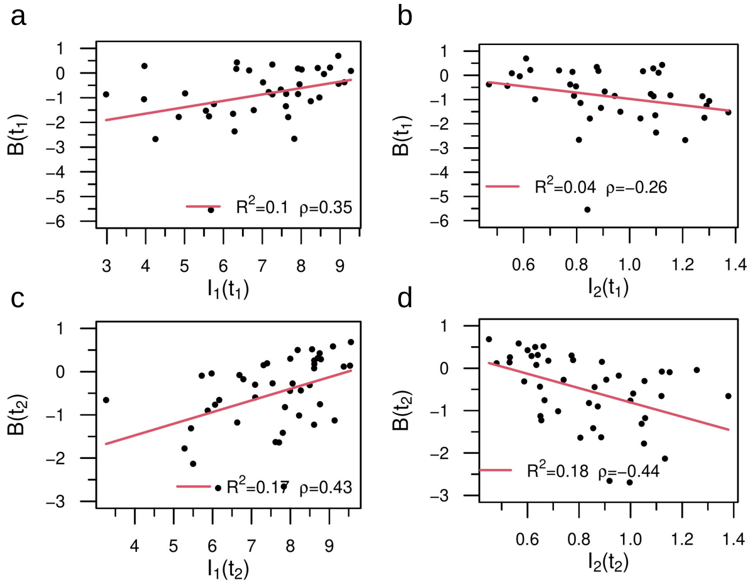


**Supplementary Figure 7** Sensitivity of the results in a uniform parcellation. Statistical correlates between dynamical variables and behavior. a,c) Correlation between $I_1$ and behavior at $t_1$ and $t_2$, respectively. The legend shows the (linear) correlation, $\rho$ and the $R^2$. b,d) Same but for $I_2$. The dynamical variables ($I_1, I_2$) predicted a significant amount of behavioral variance, higher at $t_2$. We note the very same trend (positive vs. negative correlation) but a slight deterioration in performance with respect to the Gordon's parcellation.

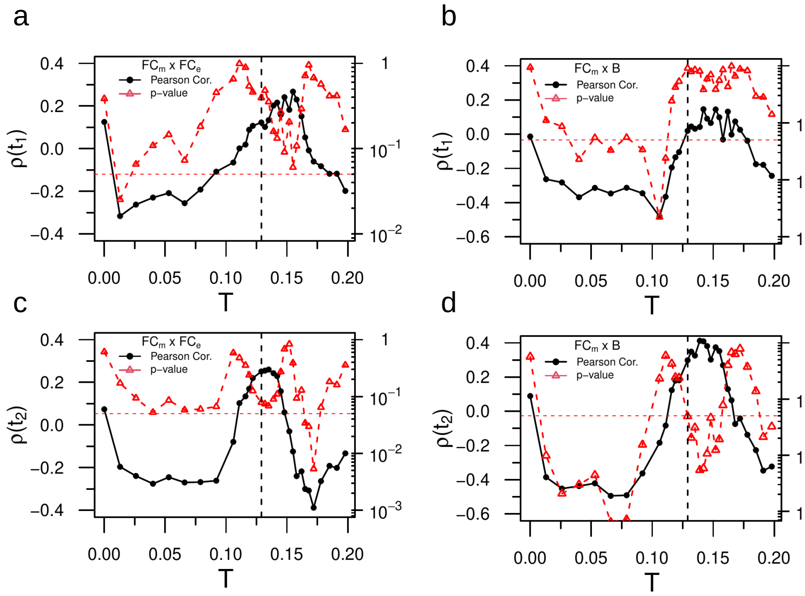


**Supplementary Figure 8** Sensitivity of the results in a uniform parcellation. Relationship between functional connectivity and behavior at criticality. a,c) Linear correlation (two-tailed) between model $FC_m$ and empirical $FC_e$ for various values of the excitation threshold $T$ for $t_1$ and $t_2$, respectively. Note the sharp increase in correlation (black dots) close to the critical point and associated behavior of the p-values (red dots). We added red horizontal lines at $p = 0.05$ for ease interpretation. b,d) The same as the previous one, but for $B$. Note the similar behavior with a peak close to the corresponding critical points (see vertical dashed line). There is a slight improvement in performance for the uniform parcellation in $t_1$, while a decrease in performance at $t_2$ compared with the Gordon's parcellation. The p-values reported in panels a-d were not corrected for multiple comparisons.

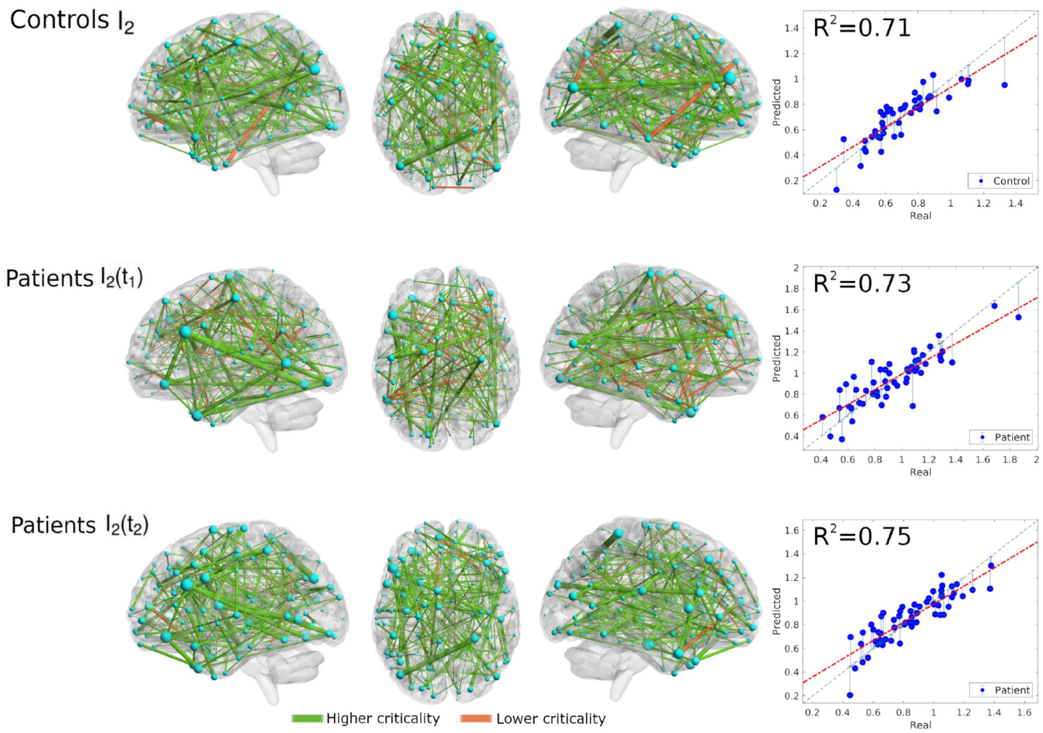


**Supplementary Figure 9** Sensitivity of the results in a uniform parcellation. Structural connectivity related to criticality (integral of the second cluster size $I_2 = \int S_2 dT$). Structural edges at $t_2$ that predict higher (green edges) and lower criticality (orange edges) values at $t_2$ (Top: controls; Middle: patients at $t_1$; Bottom: patients at $t_2$). Map of predictive connections of $W$. Only the highest 200 connections are shown for ease interpretation. The size of each ROI corresponds to the number of predictive edges converging on it. The scatter plot shows real vs. predicted criticality values from the Ridge Regression model. The gray dashed line has a slope of one and is a guide to the eye. Below we show the predictions of the Ridge Regression analyzes. For the uniform parcellation, we cannot color the brain regions according to their network membership. Both parcellations predict a large amount of variance (see $R^2$ in the scatter plots), but the uniform parcellation has slightly higher values of $R^2$. Some factors that could influence these results are: i) the higher number of regions (400 against 324) and ii) the regions are balanced in size (they all have the same size). We also show an anatomical embedding of the edges. The random parcellation lacks any physiological significance, thus limiting comparison. However, the maps of predictive edges are more balanced and points more homogeneously across the brain relative to the functional case.

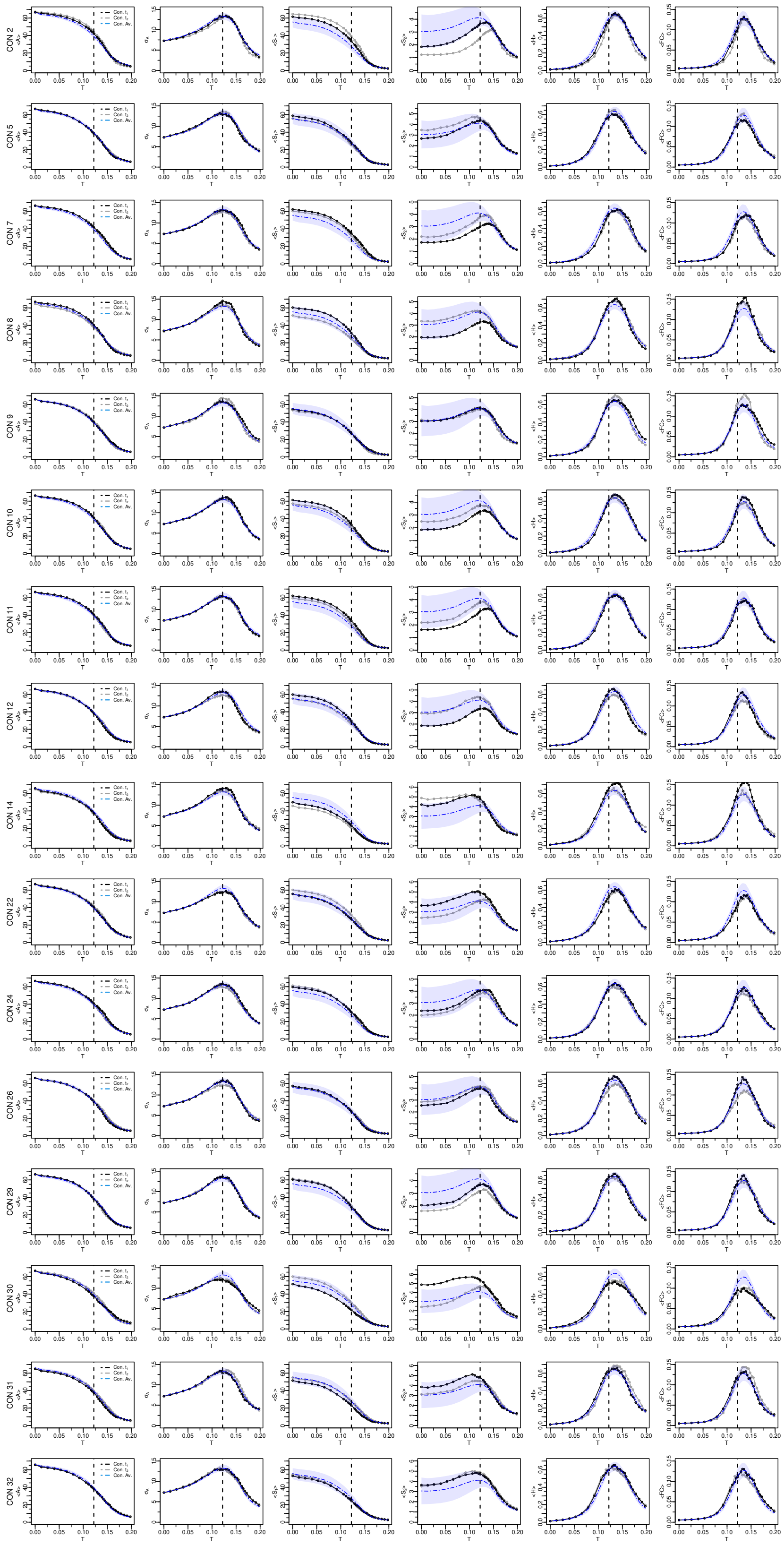

**Supplementary Figure 10** Individual neural activity patterns of healthy controls. The black lines represent controls at time $t_1$ and the gray lines at time $t_2$. In blue we show the corresponding control's group average while the shaded area is the standard deviation. From left to right: Average activity ($\langle A \rangle$), standard deviation of activity ($\sigma_A$), first cluster size ($\langle S_1 \rangle$), second cluster size ($\langle S_2 \rangle$), entropy ($\langle H \rangle$) and average functional connectivity ($\langle \mathrm{FC} \rangle$). Vertical dashed lines corresponds to the critical point ($T_c \sim 0.122$).

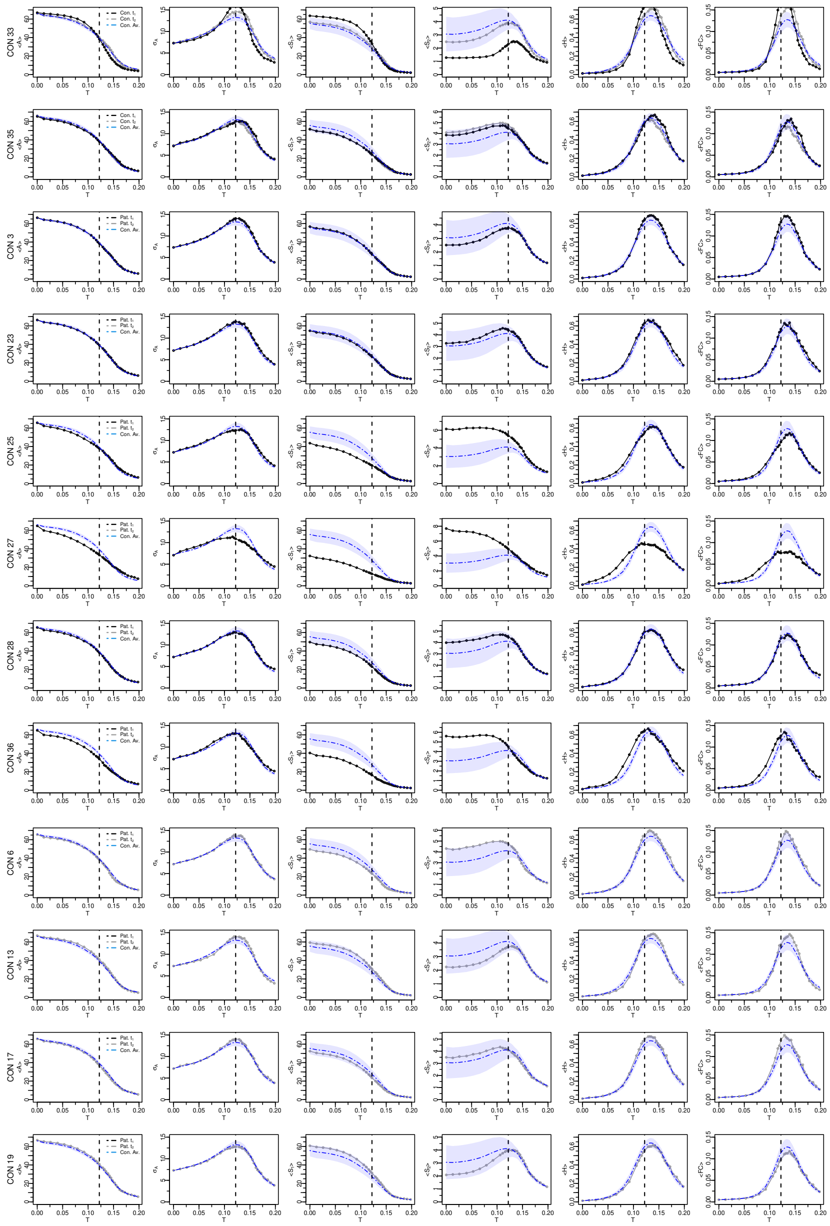

**Supplementary Figure 11** Individual neural activity patterns of healthy controls. The black lines represent controls at time $t_1$ and the gray lines at time $t_2$. In blue we show the corresponding control's group average while the shaded area is the standard deviation. From left to right: Average activity ($\langle A \rangle$), standard deviation of activity ($\sigma_A$), first cluster size ($\langle S_1 \rangle$), second cluster size ($\langle S_2 \rangle$), entropy ($\langle H \rangle$) and average functional connectivity ($\langle \mathrm{FC} \rangle$). Vertical dashed lines corresponds to the critical point ($T_c \sim 0.122$).

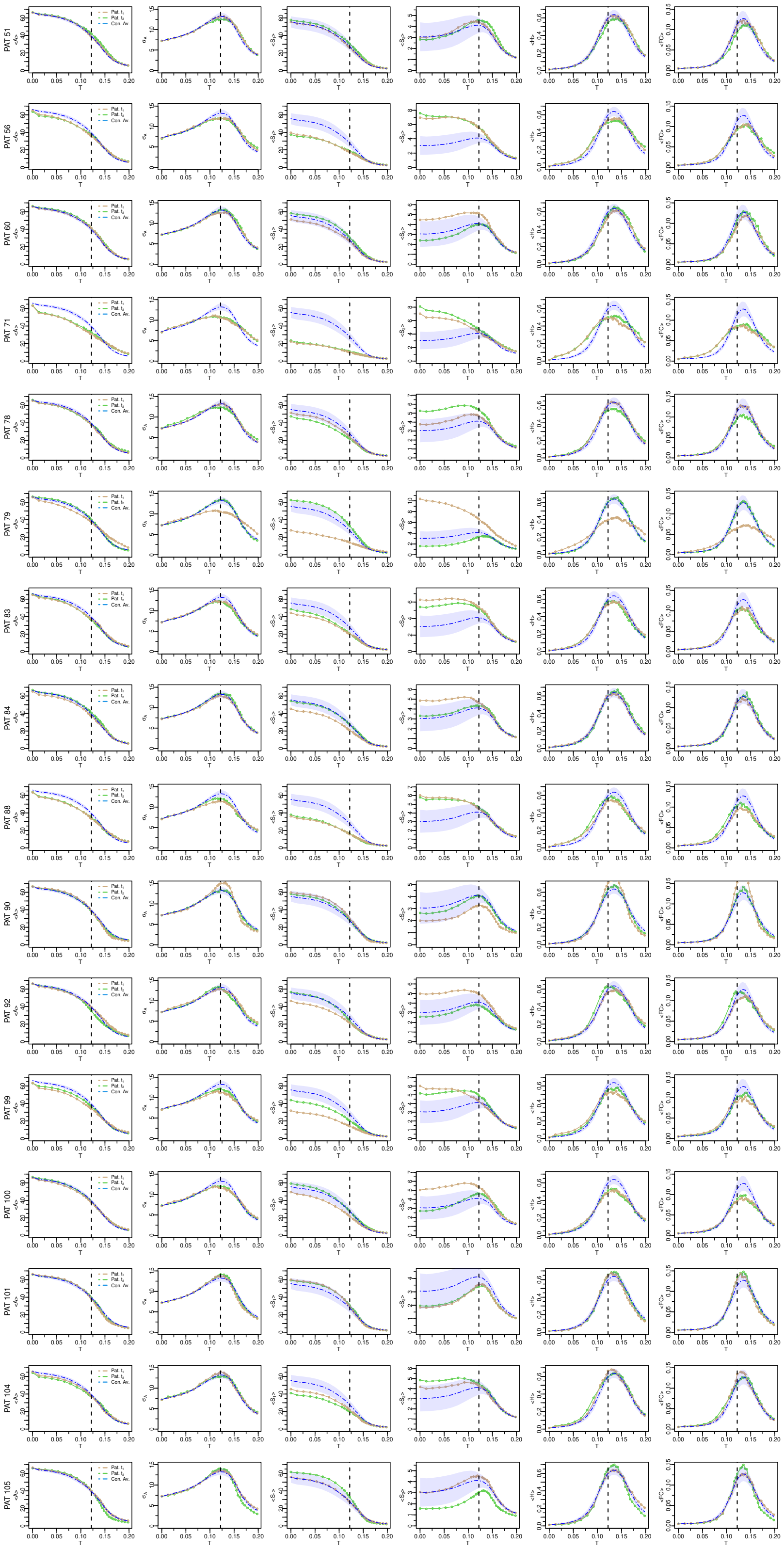

**Supplementary Figure 12** Individual neural activity patterns of stroke patients. The brown lines represent patients at time $t_1$ (3 months post-stroke) and the green lines at time $t_2$ (12 months post-stroke). In blue we show the corresponding control's group average while the shaded area is the standard deviation. From left to right: Average activity ($\langle A \rangle$), standard deviation of activity ($\sigma_A$), first cluster size ($\langle S_1 \rangle$), second cluster size ($\langle S_2 \rangle$), entropy ($\langle H \rangle$) and average functional connectivity ($\langle \mathrm{FC} \rangle$). Vertical dashed lines corresponds to the critical point ($T_c \sim 0.122$).

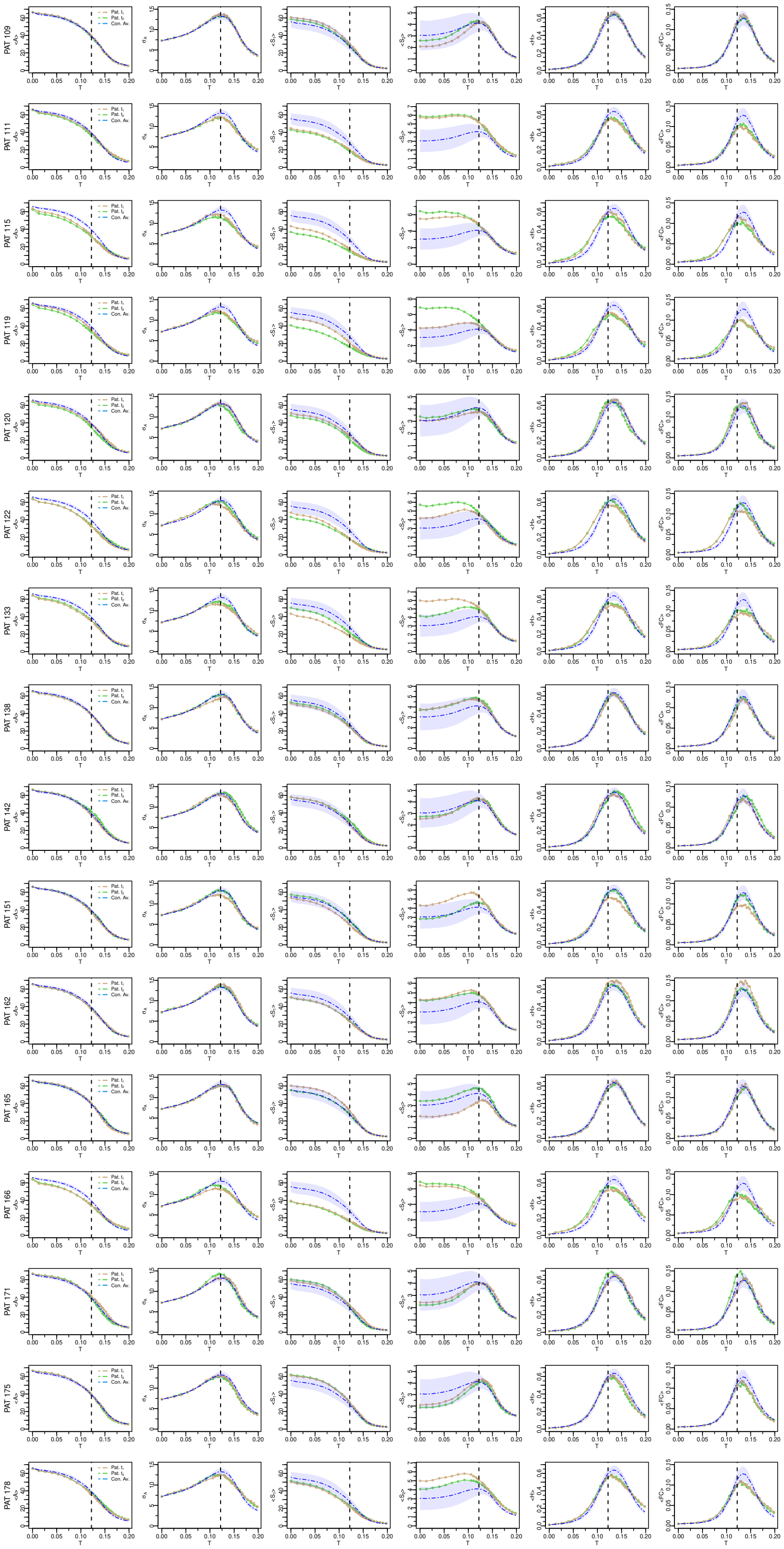


**Supplementary Figure 13** Individual neural activity patterns of stroke patients. The brown lines represent patients at time $t_1$ (3 months post-stroke) and the green lines at time $t_2$ (12 months post-stroke). In blue we show the corresponding control's group average while the shaded area is the standard deviation. From left to right: Average activity ($\langle A \rangle$), standard deviation of activity ($\sigma_A$), first cluster size ($\langle S_1 \rangle$), second cluster size ($\langle S_2 \rangle$), entropy ($\langle H \rangle$) and average functional connectivity ($\langle \mathrm{FC} \rangle$). Vertical dashed lines corresponds to the critical point ($T_c \sim 0.122$).

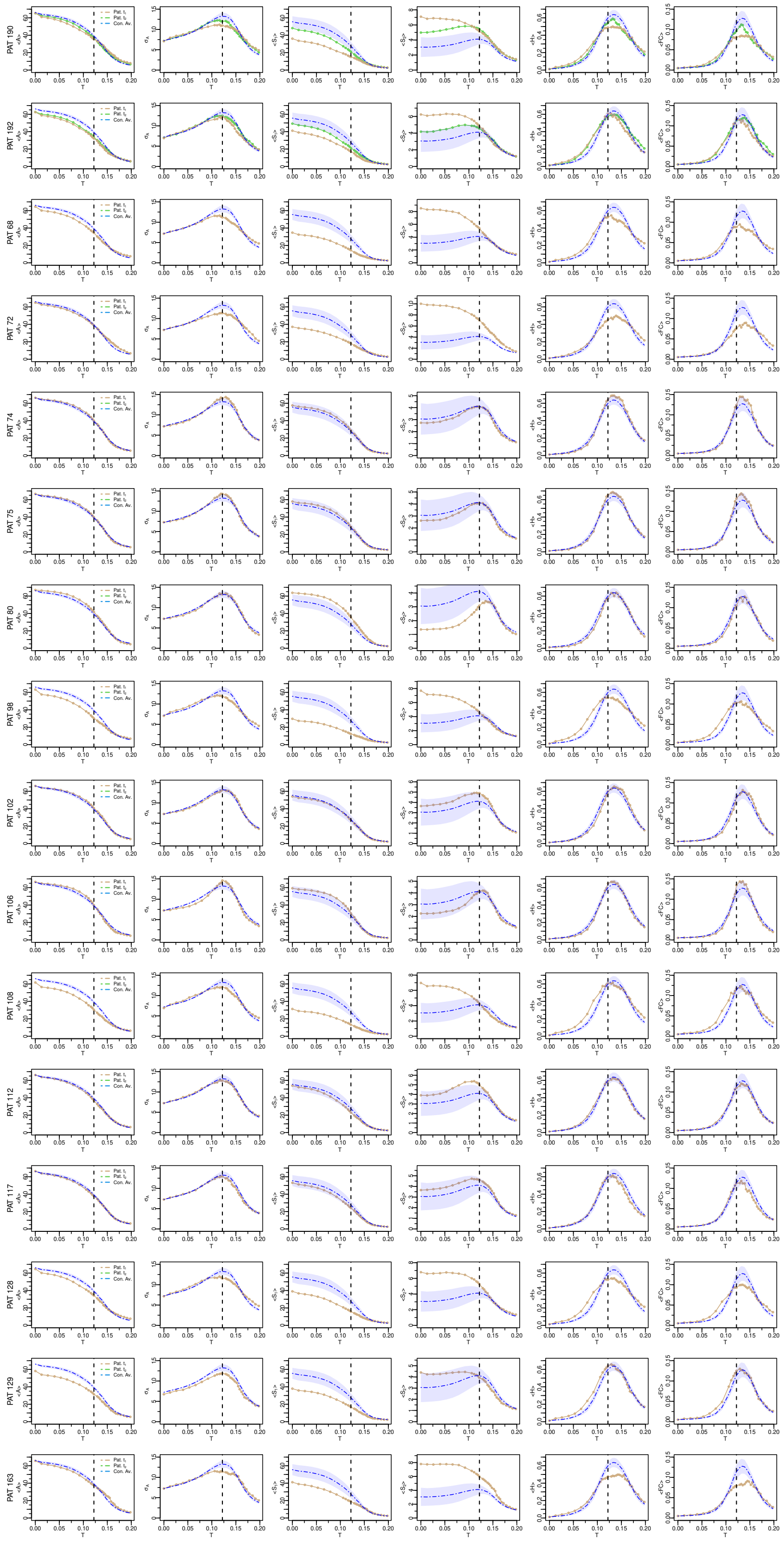


**Supplementary Figure 14** Individual neural activity patterns of stroke patients. The brown lines represent patients at time $t_1$ (3 months post-stroke) and the green lines at time $t_2$ (12 months post-stroke). In blue we show the corresponding control's group average while the shaded area is the standard deviation. From left to right: Average activity ($\langle A \rangle$), standard deviation of activity ($\sigma_A$), first cluster size ($\langle S_1 \rangle$), second cluster size ($\langle S_2 \rangle$), entropy ($\langle H \rangle$) and average functional connectivity ($\langle \mathrm{FC} \rangle$). Vertical dashed lines corresponds to the critical point ($T_c \sim 0.122$).

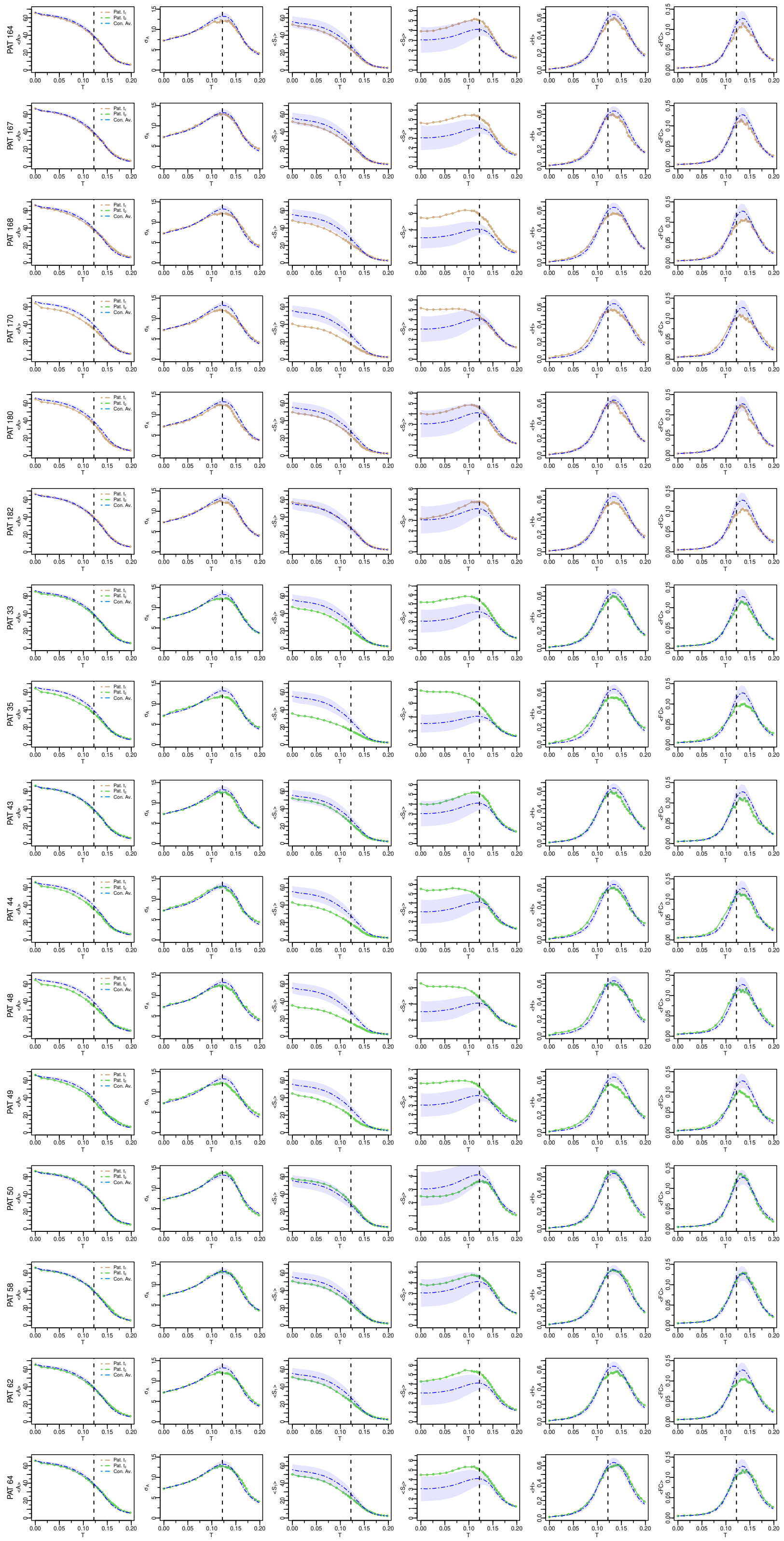


**Supplementary Figure 15** Individual neural activity patterns of stroke patients. The brown lines represent patients at time $t_1$ (3 months post-stroke) and the green lines at time $t_2$ (12 months post-stroke). In blue we show the corresponding control's group average while the shaded area is the standard deviation. From left to right: Average activity ($\langle A \rangle$), standard deviation of activity ($\sigma_A$), first cluster size ($\langle S_1 \rangle$), second cluster size ($\langle S_2 \rangle$), entropy ($\langle H \rangle$) and average functional connectivity ($\langle \mathrm{FC} \rangle$). Vertical dashed lines corresponds to the critical point ($T_c \sim 0.122$).

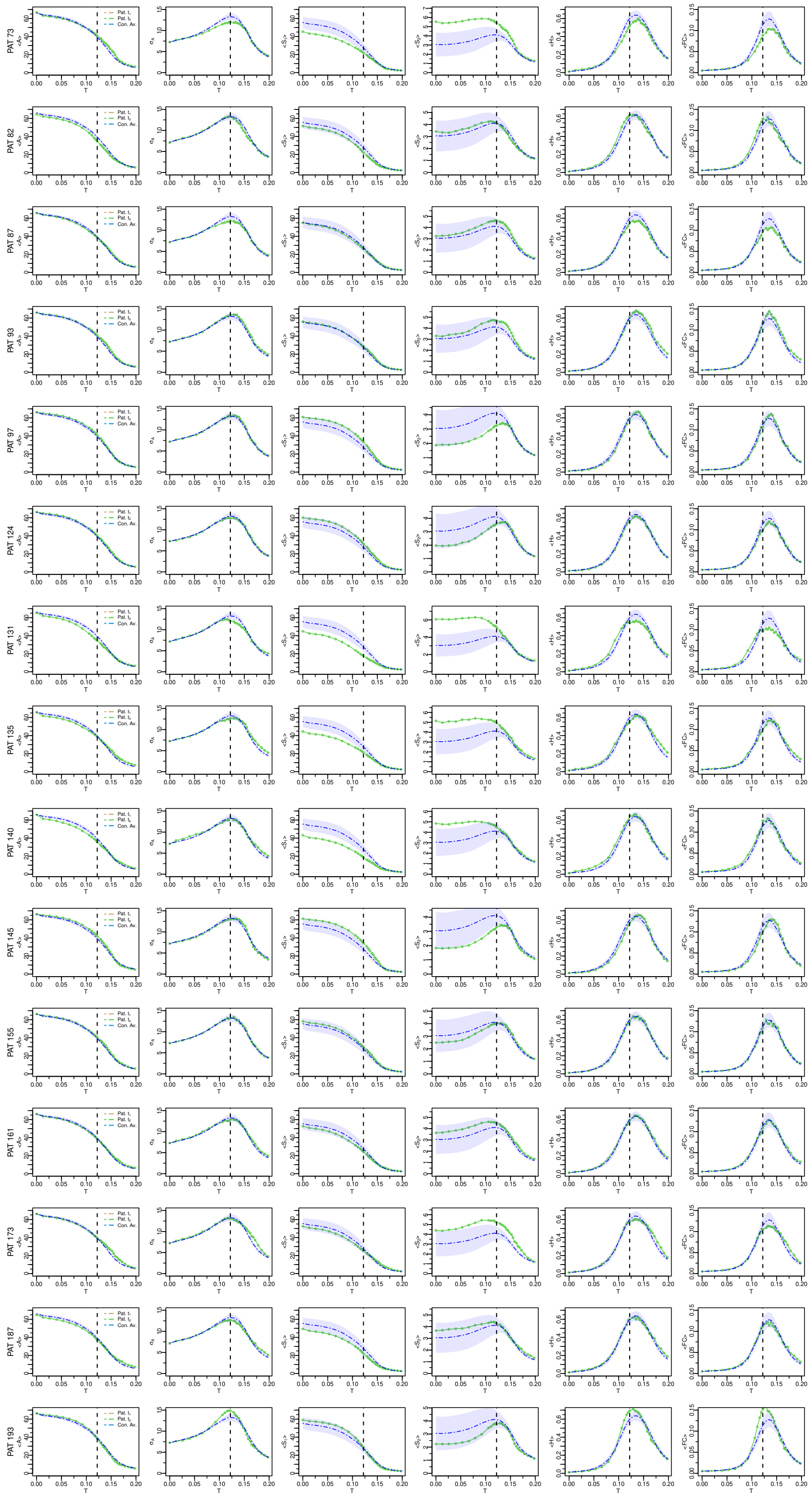


**Supplementary Figure 16** Individual neural activity patterns of stroke patients. The brown lines represent patients at time $t_1$ (3 months post-stroke) and the green lines at time $t_2$ (12 months post-stroke). In blue we show the corresponding control's group average while the shaded area is the standard deviation. From left to right: Average activity ($\langle A \rangle$), standard deviation of activity ($\sigma_A$), first cluster size ($\langle S_1 \rangle$), second cluster size ($\langle S_2 \rangle$), entropy ($\langle H \rangle$) and average functional connectivity ($\langle \mathrm{FC} \rangle$). Vertical dashed lines corresponds to the critical point ($T_c \sim 0.122$).

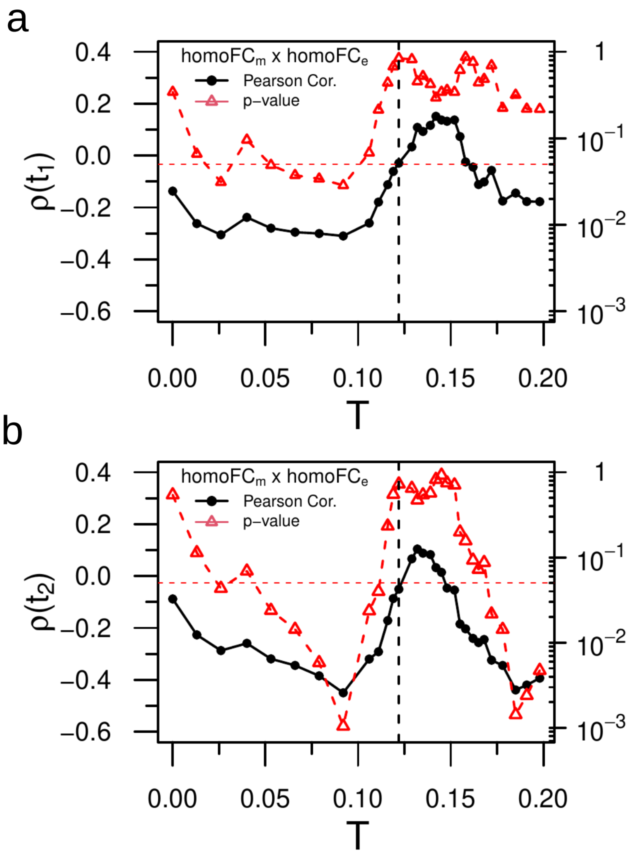


**Supplementary Figure 17** Relationship between empirical and model homotopic functional connectivity. a,b) Linear correlation (two-tailed) between model homo-FC$_m$ and empirical homo-FC$_e$ for various values of the excitation threshold $T$ for stroke patients at $t_1$ and $t_2$, respectively. We added red horizontal lines at $p = 0.05$ for ease interpretation. The p-values reported in panels a-b were not corrected for multiple comparisons.

### Supplementary Tables

**Supplementary Table 1** Alterations in network topology and associated white-matter remodeling in stroke patients from $t_1$ to $t_2$ (two-sides paired t-test). In red we show the significant p-values (FDR corrected, $\alpha = 0.05$). Two versions of the connectivity matrices were used in the analyses: weighted (based on the number of fiber tracts); binary (based on the adjacency matrix). Sample size: controls ($n = 18$); patients ($n = 34$).

| Sum of all edges (Weighted) | |
|---|---|
| Patients ($t_1$ vs. $t_2$) | Controls ($t_1$ vs. $t_2$) |
| 0.011 | 0.059 |

| Sum of all edges (Binary) | |
|---|---|
| Patients ($t_1$ vs. $t_2$) | Controls ($t_1$ vs. $t_2$) |
| 0.049 | 0.29 |

| FDR ($\alpha = 0.05$) | |
|---|---|
| Sum of all edges by RSNs networks (Weighted) | |
| Patients ($t_1$ vs. $t_2$) | Controls ($t_1$ vs. $t_2$) |
| 10 Networks FDR corrected | No effect |
| Visual | Visual |
| Retrosplenial Temporal | Retrosplenial Temporal |
| SMhand | SMhand |
| SMmouth | SMmouth |
| Auditory | Auditory |
| Cingulo Opercular | Cingulo Opercular |
| Ventral Attention | Ventral Attention |
| Salience | Salience |
| Cingulo Parietal | Cingulo Parietal |
| Dorsal Attention | Dorsal Attention |
| Fronto Parietal | Fronto Parietal |
| Default Mode | Default Mode |
| None | None |

| Sum of all edges by RSNs networks (Binary) | |
|---|---|
| Patients ($t_1$ vs. $t_2$) | Controls ($t_1$ vs. $t_2$) |
| No effect | No effect |
| Visual | Visual |
| Retrosplenial Temporal | Retrosplenial Temporal |
| SMhand | SMhand |
| SMmouth | SMmouth |
| Auditory | Auditory |
| Cingulo Opercular | Cingulo Opercular |
| Ventral Attention | Ventral Attention |
| Salience | Salience |
| Cingulo Parietal | Cingulo Parietal |
| Fronto Parietal | Fronto Parietal |
| Default Mode | Default Mode |
| None | None |

**Supplementary Table 2** Statistical correlates between dynamical, functional and behavioral patterns. In the table we show the neural variables, the linear correlation ($\rho$), the sample size ($n$), and the p-value. In red we show the significant p-values (two-tailed, FDR corrected, $\alpha = 0.05$).

| Neural variables | $n$ | $\rho$ | p-value |
|---|---|---|---|
| $B(t_1)\,, I_1(t_1)$ | 38 | 0.40 | 0.0125 |
| $B(t_2)\,, I_1(t_2)$ | 43 | 0.50 | 0.0006 |
| $B(t_1)\,, I_2(t_1)$ | 38 | -0.25 | 0.1300 |
| $B(t_2)\,, I_2(t_2)$ | 43 | -0.51 | 0.0004 |
| $B(t_1)$ , homo-FC$_e(t_1)$ | 59 | 0.43 | 0.0006 |
| $B(t_2)$ , homo-FC$_e(t_2)$ | 57 | 0.40 | 0.0019 |
| homo-FC$_e(t_1)\,, I_1(t_1)$ | 50 | 0.43 | 0.0020 |
| homo-FC$_e(t_2)\,, I_1(t_2)$ | 50 | 0.46 | 0.0008 |
| homo-FC$_e(t_1)\,, I_2(t_1)$ | 50 | -0.27 | 0.0591 |
| homo-FC$_e(t_2)\,, I_2(t_2)$ | 50 | -0.44 | 0.0014 |
| FC$_e(t_1)\,, I_1(t_1)$ | 50 | 0.23 | 0.1156 |
| FC$_e(t_2)\,, I_1(t_2)$ | 50 | 0.29 | 0.0434 |
| FC$_e(t_1)\,, I_2(t_1)$ | 50 | -0.11 | 0.4546 |
| FC$_e(t_2)\,, I_2(t_2)$ | 50 | -0.37 | 0.0098 |

**Supplementary Table 3** Analysis of variance of the neural variables between groups and across the two time points using Two-Way Mixed ANOVA. In the table we show the neural variables, the $F$ statistics, and the p-value. In the last two columns we show the planned two-tailed paired t-tets and corresponding p-values, respectively. Sample size: $n = 18$ (controls); $n = 34$ (patients).

| | Group | | Time | | Group by Time | | Patients ($t_1$ vs. t$_2$) | |
|---|---|---|---|---|---|---|---|---|
| Neural Variables | $F$ | $p$ | $F$ | $p$ | $F$ | $p$ | $t$ | $p$ |
| $S_1$ | 17.5 | 0.0001 | 1.3 | 0.25 | 2.4 | 0.12 | 2.21 | 0.03 |
| $S_2$ | 12.7 | 0.0008 | 0.003 | 0.95 | 1.2 | 0.27 | 0.96 | 0.34 |
| $A$ | 15.4 | 0.0002 | 1.9 | 0.17 | 0.05 | 0.82 | 1.27 | 0.21 |
| $\sigma_A$ | 8.8 | 0.004 | 0.1 | 0.7 | 5.5 | 0.02 | 1.57 | 0.12 |
| FC | 8.7 | 0.005 | 0.4 | 0.5 | 5.5 | 0.02 | 1.43 | 0.16 |
| $H$ | 10.6 | 0.002 | 0.26 | 0.6 | 5.9 | 0.02 | 1.50 | 0.14 |
| $I_1$ | 15.4 | 0.0003 | 1.0 | 0.3 | 2.7 | 0.10 | 2.03 | 0.05 |
| $I_2$ | 16.2 | 0.0002 | 0.4 | 0.5 | 1.6 | 0.20 | -1.45 | 0.15 |
| $K$ | 11.9 | 0.001 | 0.04 | 0.8 | 4.5 | 0.04 | 2.10 | 0.04 |
| $H_{SC}$ | 11.4 | 0.001 | 1.7 | 0.19 | 2.21 | 0.14 | 2.19 | 0.03 |
| $E$ | 15.4 | 0.0003 | 0.18 | 0.7 | 3.5 | 0.07 | 1.81 | 0.08 |
| $Q$ | 15.0 | 0.0003 | 0.004 | 0.9 | 2.2 | 0.14 | -1.21 | 0.23 |

**Supplementary Table 4** Statistical correlates between behavior ($B$) and graph topological metrics of the brain network. In the table we show the network metrics, the (linear) correlation, $\rho$ and the p-value. In red we show the significant p-values (two-tailed, FDR corrected, $\alpha = 0.05$). Sample size: $n = 38$ (time $t_1$); $n = 43$ (time $t_2$).

| Network metrics | $\rho(t_1)$ | p-value$(t_1)$ | $\rho(t_2)$ | p-value$(t_2)$ |
|---|---|---|---|---|
| $K$ | 0.3 | 0.069 | 0.54 | 0.00019 |
| $H_{SC}$ | 0.27 | 0.097 | 0.45 | 0.0026 |
| $Q$ | -0.16 | 0.33 | -0.47 | 0.0015 |
| $E$ | 0.35 | 0.034 | 0.54 | 0.00017 |

### Supplementary Note 1: White-matter remodeling

The recovery of criticality from three to twelve months must reflect a change in the underlying structural connectivity. We performed two analyses with different approaches to investigate the degree of the network reorganization. First, a very general one in which we checked the difference between $t_1$ and $t_2$ of the whole matrix as a sum of all edges (1 comparison only). This approach tests whether the SC matrices, or portions of them, have changed over time in terms of the number of white-matter fibers. Second, a much more specific one in which the sum was done per node (324 comparisons). For both approaches we used the DWI structural connectivity (number of fiber tracts) and the corresponding adjacency matrices. Therefore, in addition to a remodeling of white-matter connections, we tested whether there was also a difference in terms of emergence of new white-matter connections, and therefore a change in topology. In the first case, we find significant difference (paired t-test) in patients in both DWI and adjacency (topology) matrices (Supplementary Table 1). In the second case, however, differences emerged but were not confirmed after FDR correction (results not shown). This is explained on the one hand by too many comparisons, and on the other by the high variability between patients, which does not allow us to find a systematic change over time at the level of a single node. We therefore proposed a halfway approach between the two and did the same statistics at brain network level. Specifically, we summed up all the edges per network (13 comparisons). A very clear difference emerged in the SC matrices of patients for 10 out of 13 networks while no effect emerged for topology (Supplementary Table 1). These results shows that at a global level, in the brain, we have changes in both the number of fibers and topology, while at the level of individual networks there is more evidence of remodeling of white-matter connections (increasing fibers in existing connections) rather than recovery of new white-matter connections. We also performed all these analyses for the controls and found no differences, as expected.

### Supplementary Note 2: Behavioral battery

The present stroke cohort has been investigated with a large battery of neurobehavioral tests. The battery includes most of the tests recommended by the National Institute of Neurological Disorders and the Canadian Stroke Network task force for the harmonization of cognitive impairment measures in vascular diseases[1]. The behavioral battery also includes tests of body function according to the ICF classification. We also use a computerized (Posner) task to measure attention processes that has been carefully characterized in terms of sensitivity and specificity[2]. Below we describe the tests used (for additional information see Corbetta et al.[3] and the references therein).

#### Motor Battery

Upper body function was measured in both arms as follows:

1. Active range of motion against gravity measured by goniometry at Shoulder Flexion, and Wrist Extension;

2. Grip strength measured by dynamometry;

3. Dexterity measured with the 9-Hole Peg Test in which patients placed nine plastic pegs into holes on a pegboard as quickly as possible (pegs/second);

4. Function measured with the Action Research Arm Test total score (ARAT), in which patients performed = functional grasp, grip, pinch, and gross motor movements according to the standardized protocol and were rated for quality of movement;

5. Combined Walking Index: Patients were timed while walking 10 meters if able to safely do so unassisted. Patients who were unable to walk 10 meters were rated using the Walking item on the Functional Independence Measure;

6. Left/right Total Motricity Index (MI), which sums the manual muscle testing scores for left/right hip flexion, knee extension, ankle dorsiflexion;

7. Ankle dorsiflexion goniometry for left/right active range of motion against gravity.

#### Language Battery

Subtests of the Boston Diagnostic Aphasia Examination (BDAE-III) were performed according to the standard protocol:

1. Basic word discrimination: Patients pointed to the picture that matched the word named by the experimenter;

2. Commands: Patients performed one- to five-step commands spoken by the experimenter;

3. Complex Ideational Material: Patients answered yes/no questions spoken by the experimenter;

4. Boston Naming Test short form: Patients named the item pictured;

5. Oral Reading of Sentences: Patients read sentences aloud;

6. Comprehension of Oral Reading of Sentences: Patient answered multiple-choice comprehension questions about the sentences they just read; In addition we measured phonetic/phonological processing with:

7. Non-word Reading: Four-letter nonwords (e.g. NORD) were presented. Patients were instructed to say the nonword aloud;

8. Stem Completion: Three-letter word stems (e.g., COU) were presented. Patients were instructed to say a word that started with those three letters (e.g., COUPLE). This task provides a very stereotypical pattern of language-related areas.

#### Executive Function

A subtest of the Delis-Kaplan Executive Function System was performed according to the standard protocol: Animal Naming: Patients named as many animals as possible in 1 minute.

### Memory Battery

Visual memory was studied with the Brief Visuospatial Memory Test-Revised (BVMT-R). Patients studied abstract figures and were asked to reproduce them from memory on three immediate recall trials and one delayed recall trial. After the delayed recall trial, patients were shown figures and asked if each was one of the studied figures. Scores were calculated for the following variables:

1. BVMT Immediate Total Recall T-score: age-normed using the tables provided in the test manual;
2. BVMT Delayed Recall T-score: age-normed using the tables provided in the test manual;
3. BVMT Delayed Recall percent retained: calculated from the percent items retained from last immediate recall trial to delayed recall;
4. BVMT Delayed Recognition discrimination index: calculated from proportion of correct recognitions, correct rejections, misses, and false alarms, using the table provided in the test manual.

Verbal memory was assessed with the Hopkins Verbal Learning Test-Revised (HVLT-R). Patients listened to a list of words and were asked to repeat them from memory on three immediate recall trials and one delayed recall trial. After the delayed recall trial, patients were read a list of words and asked if each was one of the studied words. Scores were calculated for the following variables:

1. HVLT Immediate Total Recall T-score: age-normed using the tables provided in the test manual;
2. HVLT Delayed Recall T-score: age-normed using the tables provided in the test manual;
3. HVLT Delayed Recall percent retained: calculated from the percent items retained from last immediate recall trial to delayed recall;
4. HVLT Delayed Recognition discrimination index: calculated from the proportion of correct recognitions, correct rejections, misses, and false alarms, age-normed using the table provided in the test manual.

Spatial working memory was examined with a subtest of the Wechsler Memory Scale:

Spatial Span: Patients watched the examiner tap sequences on a block board and then asked to copy the sequences. After reaching a performance ceiling, the patients watched the examiner tap sequences on the block board and then asked to produce the sequences in reverse order. Scores were calculated for the following variables: 1. Spatial Span Forwards; 2. Spatial Span Backwards

### Attention Battery

Different visuospatial attention processes were measured with the Posner orienting task. The test took a total of 15 minutes to administer, including a practice block. The following scores were calculated:

1. Visuospatial contralesional biases were measured with:

   (a) The Visual Field Reaction Times (RT): relative delay in RTs for targets presented in the left vs. right visual field

   (b) The Visual Field Accuracy: relative percent misses for targets presented in the left vs. right visual field

2. Deficits in shifting attention were measured with:

   (a) The Validity Effect RT: relative delay in RTs for targets presented following valid vs. invalid cues

   (b) The Validity Effect Accuracy: relative percent misses for targets presented following valid vs. invalid cues

3. Deficits in re-orienting to unattended locations with:

   (a) The Disengagement Effect RT: relative delay in RTs for targets presented in the left visual field following an invalid cue.

   (b) The Disengagement Effect Accuracy: relative percent misses for targets presented in the left visual field following an invalid cue.

4. Overall performance or sustained attention were measured with:

   (a) Average RT: average of the RTs across all four conditions.

   (b) Average accuracy: average percent misses across all four conditions.

Visuomotor spatial deficits were assessed with the Star Cancellation subtest of the Behavioral Inattention Test (BIT), and the Mesulam Unstructured Symbol Cancellation Test. The following score were calculated:

1. Mesulam Center-of-cancellation, L-R misses: which reflects the lateralized center of mass of misses, using the software provided by Rorden and Karnath, for left-sided vs. right-sided misses.

2. BIT: star cancellation, Center-of-cancellation, L-R misses: which reflects the lateralized center of mass of hits, using the software provided by Rorden and Karnath, for left-sided vs. right-sided misses.

### Supplementary Note 3: Correlation between network topology and behavior

We provide an additional compelling evidence of the intricate relationship between brain criticality, the underlying network topology and behavior. Given the average degree ($K$) and the connectivity disorder ($H_{SC}$) are responsible for the underlying criticality regime alterations[4], then we hypothesize that these same variables should predict the patients behavioral performance as well. Remarkably, we verified this theoretical assumption in our empirical dataset of stroke (Supplementary Table 4). Beyond that, we find the same normalization patterns across time-points as described previously, thus suggesting the recover of the normal structure-function (behavior) relationship. Indeed, the network metrics predicted a large amount of behavioural variance at $t_2$. Apart from the global efficiency (E), these same variables did not show statistically significant correlations at $t_1$.

### Supplementary Note 4: Individual model neural analysis

In the Supplementary Figures 10-16 we show the individual neural profiles for healthy controls and stroke patients. Each row contains data from one subject, either control (CON ID - age-matched control)) or stroke patient (PAT ID - patient's identifier). The thin solid curves represent individual data while bold curves represent the group average, i.e., $X^{av} \equiv \sum_i^n X(i)/n$, where $n$ is the number of individuals in each group; shaded areas correspond to one standard deviation of the group means. We show the following neural variables: the average activity, $\langle A \rangle$; the standard deviation of the activity, $\sigma_A$; the average of the first cluster size, $\langle S_1 \rangle$; the average of the second largest cluster, $\langle S_2 \rangle$; the average entropy, $\langle H \rangle$; and the average functional connectivity, $\langle \mathrm{FC} \rangle$.

In order to follow the temporal evolution of the neural variables, we plotted the model predictions for the stroke patients for two time points, such as $t_1$ (3 months post-stroke) and $t_2$ (12 months post-stroke). Since not all individuals have participated in the two scan sections, in a few cases data is missing for $t_1$ or $t_2$. The control's group average is also illustrated in blue solid line.

To facilitate interpretation, we describe briefly the general behavior of the simulated variables as a function of $T$. In our model, $T$ plays the role of an inhibition parameter that regulates the amount of input excitatory activity. Therefore, the super-critical phase ($T \ll T_c$), is characterized by high excitation (high $\langle A \rangle$), that in turn is dominated by a giant cluster size (high $S_1$ and low $S_2$). However, these fluctuations are uncorrelated (low FC) and the complexity of the functional connections is very low (low $H$). On the other side, high values of $T$ ($T \gg T_c$), represents a system with a lot of inhibition (sub-critical phase). The average activity approaches a small constant value, with decreased values of $\sigma_A$, $S_2$, $H$, and FC. Finally, at the critical point ($T = T_c$), we observe a significant increase in neural fluctuations, $\sigma_A$, entropy, and long-range correlations (high FC).

Now, we turn our attention to the controls data (Supplementary Figures 10-11). A careful visual inspection reveals that intra- and inter-subject variability are very small; in other words, the neural patterns are well behaved. However, few exceptions do exists, for instance, the individual's $\mathrm{ID} = \{25, 27, 36\}$. This variation from the control's baseline may be related to inherent features of the brain's anatomy, and/or experimental fluctuations/errors in the data acquisition.

Now we consider the patient's group (Supplementary Figures 12-16). In contrast with the previous case, we observe a great variability in neural patterns in stroke, either across individuals as well as across time points. However, in general, there is a normalization pattern across time points, i.e., the neural variables approaches the control's average at 12 months. Note that the second largest cluster, $S_2$, decreases with time. This reflects a normalization in the integration-segregation balance. In $t_1$, activity clusters are more segregated, as expressed by large $S_2$ (or correspondingly, small $S_1$, i.e., the giant component is broken down into smaller clusters). With recovery, such a balance is normalized, and $S_2$ approaches the controls baseline, indicating a more integrated dynamics.

### Supplementary Note 5: Uniform parcellation

In this section we investigate the sensitivity of the results with respect to the brain parcellation. Indeed, the effect of brain parcellation in whole-brain models has been largely neglected in previous studies (but see a recent study by Jung et al.[5]).

To fill this gap, we test the robustness of our results by replicating our analyses in a uniform parcellation[6]. Specifically, we characterized each brain voxel uniquely on the basis of its anatomical location, in terms of $x$, $y$, and $z$ coordinates. The matrix of anatomical coordinates of the brain was then fed into the k-means clustering in python. We asked the algorithm to parcellate the brain in 400 clusters, to enable the comparison with the Gordon parcellation. The uniform parcellation we employed, all the ROIs have the same size but their locations in the brain do not have any physiological or anatomical significance.

In the Supplementary Figures 5-9 we show the model predictions for the uniform parcellation. Some results from the main text are not exactly reproducible due to the lack of the RSNs network assignment or the concept of homotopy. A rigorous comparison between the two sets of results (uniform versus functional) shows that the uniform parcellation reproduces qualitatively and quantitatively all the results obtained with the Gordon's parcellation. In other words, the brain parcellation has little effect on the modeled neural dynamics. These results are quite interesting from a physics perspective. Indeed, we demonstrate that the critical features of the model are insensitive to the microscopic details of the brain parcellation, which is consistent with the theory of critical phenomena. Although we use ”uniform parcellation”, the structural matrix still keeps biological significance in the sense it correctly captures the brain connectivity.
